# Contrasting patterns of extrasynaptic NMDAR-GluN2B expression in macaque subgenual cingulate and dorsolateral prefrontal cortices

**DOI:** 10.1101/2025.02.05.636752

**Authors:** MKP Joyce, D Datta, J Arellano, A Duque, YM Morozov, JH Morrison, AFT Arnsten

## Abstract

Expression of the N-methyl-D-aspartate receptor, particularly when containing the GluN2B subunit (NMDAR-GluN2B) varies across the prefrontal cortex (PFC). In humans, the subgenual cingulate cortex (SGC) contains among the highest levels of NMDAR-GluN2B expression, while the dorsolateral prefrontal cortex (dlPFC) exhibits a more moderate level of NMDAR-GluN2B expression. NMDAR-GluN2B are commonly associated with ionotropic synaptic function and plasticity, and are essential to the neurotransmission underlying working memory in the macaque dlPFC in the layer III circuits afflicted in schizophrenia. However, NMDAR-GluN2B can also be found at extrasynaptic sites, where they may trigger distinct events, including some linked to neurodegenerative processes. The SGC is an early site of tau pathology in sporadic Alzheimer’s Disease (sAD), which mirrors its high NMDAR-GluN2B expression. Additionally, the SGC is hyperactive in depression, which is treated with NMDAR antagonists. Given the clinical relevance of NMDAR in the SGC and dlPFC, the current study used immunoelectron microscopy (immunoEM) to quantitatively compare the synaptic and extrasynaptic expression patterns of NMDAR-GluN2B across excitatory and inhibitory neuron dendrites in the rhesus macaque SGC and dlPFC. We found a larger population of extrasynaptic NMDAR-GluN2B in dendritic shafts and spines of putative pyramidal neurons in SGC as compared to the dlPFC, while the dlPFC had a higher proportion of synaptic NMDAR-GluN2B. In contrast, in putative inhibitory dendrites from both areas, extrasynaptic expression of NMDAR-GluN2B was far more frequently observed over synaptic expression. These findings may provide insight into varying cortical vulnerability to alterations in excitability and to neurodegenerative forces.

**Scope Statement:** NMDAR are ionotropic receptors that contribute to neurotransmission and second messenger signaling events. NMDAR can induce a diverse array of neuronal events, in part due to variation in subunit composition and subcellular localization of receptor expression. Expression of the GluN2B subunit varies across the prefrontal cortex in humans. This subunit is highly expressed in the subgenual cingulate, an area associated with mood and emotion, and more moderately expressed in the dorsolateral prefrontal cortex, an area associated with cognitive processes. Extrasynaptic NMDAR, which often contain with the GluN2B subunit, have been linked to detrimental cellular events like neurodegeneration. Here, using high resolution electron microscopy in rhesus macaques, we found evidence that extrasynaptic NMDAR-GluN2B expression may be more prominent in subgenual cortex than in the dorsolateral prefrontal cortex. Conversely, synaptic NMDAR-GluN2B may be more prominent in the dorsolateral prefrontal cortex, consistent with their essential contribution to neuronal firing during working memory. These findings may help to illuminate the propensity of the subgenual cortex to tonic hyperactivity in major depression and its vulnerability to neurodegeneration in Alzheimer’s disease, and may help to explain how rapid acting antidepressants exert therapeutic action across diverse neural circuits.

## Introduction

The NMDAR is a di-or tri-heteromeric complex that typically contains at least one NR1 subunit and a combination of NR2 or NR3 subunits, which define the biophysical properties of the receptor (Paoletti et al., 2013; Wyllie et al., 2013). The GluN2B subunit (encoded by the gene *GRIN2B*) confers slow closing kinetics and high calcium permeability, thus allowing significant calcium entry into the neuron (Erreger et al., 2005; Glasgow et al., 2015). In humans, *GRIN2B* expression increases along a sensory-association cortical gradient which is correlated with cortical hierarchy (Burt et al., 2018), and similar to a cortical gradient of increasing intrinsic timescales for local processing (Murray et al., 2014). *GRIN2B* expression is thus lowest in sensory areas like primary visual and somatosensory cortex, intermediate levels in association cortices such as the dorsolateral prefrontal cortex (dlPFC), and highest in limbic areas like the subgenual cingulate cortex (SGC) (Burt et al., 2018; Yang et al., 2018), an area associated with mood, visceromotor function, and hyperactivity in depression (Mayberg et al., 2005; Alexander et al., 2019a). *GRIN2B* also increases within the frontal pole across primate phylogeny, suggesting a prominent role in higher cognition (Muntané et al., 2015). Studies in macaque dlPFC suggest that the long calcium influx conferred by the NMDAR-GluN2B subunit in layer III dlPFC pyramidal cell recurrent circuits is critical to maintain persistent firing in the absence of external sensory stimulation during the delay epoch of a spatial working memory task (Lisman et al., 1998; Wang et al., 2013a; Yang et al., 2018; Yang et al., 2021). Consistent with the critical role of NMDAR-GluN2B in delay-related firing in dlPFC, post-embedding immunoEM has detected prominent NMDAR-GluN2B expression within the post-synaptic density (PSD) in spines of layer III (Wang et al., 2013a). Higher expression of NMDAR-GluN2B in limbic association areas may support neural constructs that require longer continuity than working memory, such as mood and emotion (Yang et al., 2021).

NMDAR can also be extrasynaptic where they may serve distinct functions (Petralia, 2012; Groc and Choquet, 2020; Petit-Pedrol and Groc, 2021). The presence of the GluN2B subunit has been prominently associated with extrasynaptic expression (Tovar and Westbrook, 1999; Papouin et al., 2012), though other evidence suggests that extrasynaptic subunit specificity is not quite so clear cut (Thomas et al., 2006; Harris and Pettit, 2007; Kortus et al., 2023), and that expression patterns could be area- or species-specific (e.g., (Harris and Pettit, 2007; Petralia et al., 2010; Wang et al., 2013a)). Extrasynaptic NMDAR have been associated with events distinct from synaptic NMDAR [reviewed in (Hardingham and Bading, 2010; Gladding and Raymond, 2011; Parsons and Raymond, 2014)], and in particular with a host of very detrimental events in states of acute injury, cellular distress, and neurodegeneration (e.g., (Hardingham et al., 2002; Zong et al., 2022); reviewed in (Hardingham and Bading, 2010; Gladding and Raymond, 2011; Parsons and Raymond, 2014)]).

Memantine, a low affinity, non-competitive antagonist and open channel blocker of the NMDAR, is an FDA-approved drug with modest efficacy in treating moderate-to-severe sAD (Kim et al., 2024), and is hypothesized to target extrasynaptic NMDAR (Xia et al., 2010). Further, other non-specific NMDA antagonists, like ketamine, and GluN2B-specific antagonists may target extrasynaptic populations of NMDAR to exert rapid-acting antidepressant effects (Miller et al., 2014; Miller et al., 2016; Brown and Gould, 2024; Krystal et al., 2024), though ketamine can also mimic schizophrenia by impairing dlPFC cognitive function (Beck et al., 2020). Given the significance of NMDAR-GluN2B to dlPFC and SGC function, as well as its relevance in psychiatric disorders (Yang et al., 2021), it is critical to understand the distribution of NMDAR-GluN2B across synaptic and extrasynaptic membrane domains of diverse cell types in these circuits. Variability in NMDAR membrane expression across the PFC landscape may produce mixed effects during systemic administration of pharmacological NMDAR agents.

Very few studies have examined NMDAR-GluN2B in primate PFC. As mentioned above, post-embedding immunoelectron microscopy (immunoEM) demonstrated NMDAR-GluN2B within glutamate-like synapses on spines in layer III dlPFC (Wang et al., 2013a). Post-embedding immunoEM is ideal for identifying synaptic proteins because it provides superior access to the PSD (Petralia and Wang, 2021). For example, Wang et al. (2013a) showed prominent NMDAR-GluN2B labeling within the PSD, as illustrated in Figure 1. However, the post-embedding process degrades extra-synaptic membranes, and thus the selective labeling of the PSD in Wang et al. (2013a) may be an accurate reflection of GluN2B selective localization, or may be impacted by inherent constraints of the technique. Pre-embedding immunoEM, on the other hand, often provides superior preservation of ultrastructure, in particular extrasynaptic membranes, and intracellular organelles, for example the spine apparatus, which interacts with receptors in the membrane in dlPFC for calcium-mediated calcium release (Arnsten et al., 2021b; Datta et al., 2024). Pre-embedding immunoEM, however, likely undersamples from the PSD (Petralia and Wang, 2021). Here, we have used pre-embedding immunoEM and quantitative analyses to compare the distribution of NMDAR-GluN2B membrane expression across the macaque SGC and dlPFC.

**Figure 1.**
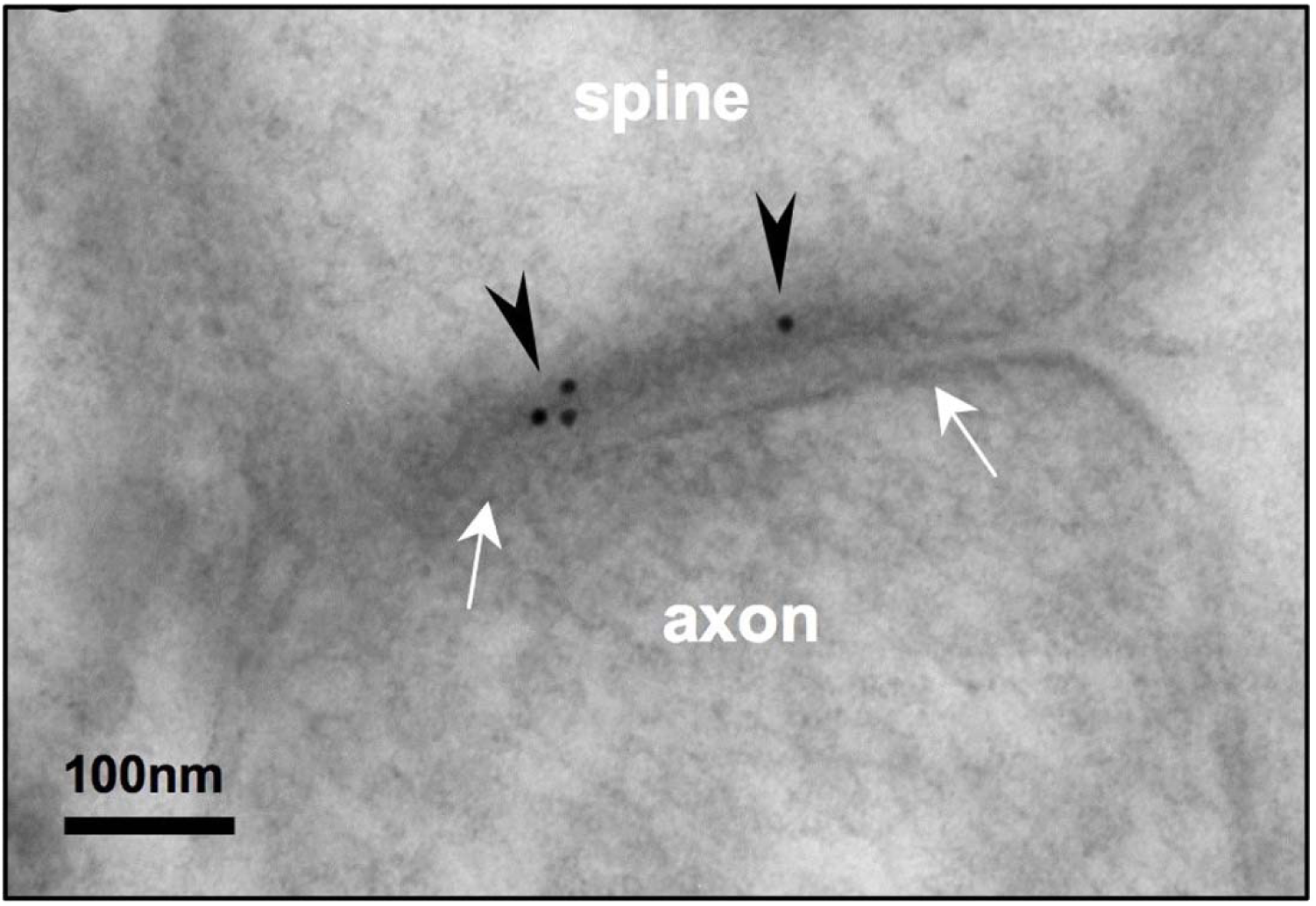
Post-embedding immunoEM for NMDA-GluN2B reveals labeling in the postsynaptic density of spines of layer III rhesus macaque dlPFC. Electron micrograph depicting a layer III dlPFC spine receiving a synapse (white arrows) from an axonal bouton. Post-embedding immunoEM preparation reveals immunogold particles (black arrowheads) labeling NMDAR-GluN2B in the post-synaptic density. Adapted from Wang et al. (2013a).

## Methods

### Experimental Design

This study was designed to assess the subcellular location of NMDAR-GluN2B in pyramidal and inhibitory neurons of layer III in dlPFC and SGC. Using pre-embedding immunoEM, we have labeled the NMDAR-GluN2B with immunogold particles in dlPFC and SGC. We have cut ultrathin sections, and we have performed systematic sampling using high resolution electron microscopy within the antibody penetration zone. We analyzed all immungold particles to determine the identity of their parent structure, and their location within that structure, to determine if there were detectable differences in the membrane distribution of these particles between the dlPFC and SGC. To determine if there were differences in NMDAR-GluN2B expression across types of inhibitory neurons, we performed multi-label immunofluorescence (MLIF) for the calcium-binding proteins (CBP) parvalbumin (PV), calbindin (CB), and calretinin (CR), in addition to NMDAR-GluN2B, and analyzed the NMDAR-GluN2B expression across CBP type using Qupath.

### Subjects and tissue selection

Research was conducted with the approval of the Yale University IACUC under NIH and USDA guidelines. Tissue from the dlPFC and SGC blocks of two young adult (aged 8 and 10years, female) macaques were used in this study. Subjects were deeply anesthetized and underwent transcardial perfusion with 0.1M phosphate-buffered saline (PBS), followed by 4% paraformaldehyde and 0.05% glutaraldehyde in 0.1M PBS. Brains were removed, and cut into blocks, including the dlPFC and SGC. Blocks were cut on a vibratome (Leica, Norcross GA USA) at a thickness of 60 µm. Free-floating sections underwent cryoprotection in ascending concentrations of sucrose (JT Baker, cat #4072-01) solution (10, 20, 30% in PBS, each overnight), and then rapidly frozen in liquid nitrogen for long-term storage at −80°C. Sections were selected from the dlPFC block along the principal sulcus, for the most part anterior to the start of the arcuate sulcus. Tissue was selected from the SGC block along the anterior to posterior breadth of A25, as long as the corpus callosum was in plane.

### Primary Antibodies used for Immunohistochemistry

To label NMDAR-GluN2B, we used the Alomone polyclonal rabbit anti-NMDAR2B antibody at 1:100 (Alomone, cat# AGC- 003, RRID:AB_2040028). This antibody has been used for MLIF in various tissues, such as cultured rat neurons (Schweitzer et al., 2017), human-induced pluripotent stem cell-derived neurons (Telezhkin et al., 2016), and *in situ* macaque dlPFC (Datta et al., 2024). This antibody binds to an antigen site corresponding to amino acid residues 323-337 of the rat NMDA receptor 2B on the extracellular N-terminus. While an antibody test in conditional or full NMDA-GluN2B knockout animals is preferable as a negative control, we were unsuccessful in locating such a test using this antibody. We performed a preadsorption control using the Alomone NMDAR-GluN2B blocking peptide (cat #BLP-GC003, <u>Fig. S6A-B</u>), and observed negligible labeling, suggesting that the regions outside the paratope of the antibody have minimal interactions in our tissue. In the dlPFC and SGC of each case, we further assessed the specificity of antibody labeling by measuring the percent of immunogold particles found inside mitochondria, which we deemed as non-specific labeling. This percent of immunogold particles found within mitochondrial boundaries ranged from 0.2-0.7%. This was much smaller than the surface area per section occupied by mitochondria, which ranged from 6-8%, suggesting that stochastic binding of the antibody was quite low.

To label the CBPs, we used Swant mouse monoclonal antibodies at 1:2000 (Swant guinea pig anti-PV cat# GP72 RRID:AB_2665495, mouse anti-CB-D28k cat# 300 RRID:AB_10000347, and goat anti-calretinin cat# CG1 RRID: AB_10000342, respectively), which have been widely used in macaque tissue (e.g., (Joyce et al., 2020; Tsolias and Medalla, 2021; Medalla et al., 2023)). To label microtubule-associated protein-2 (MAP2), we used a chicken anti-MAP2 at 1:1000 (Abcam, cat# AB5392), which has been used previously in macaque tissue as well (Tsolias and Medalla, 2021).

### Immunohistochemical Procedures for Electron Microscopy

#### Single label gold immunoEM for NMDAR-GluN2B

Free-floating sections from dlPFC and SGC underwent antigen retrieval in 10nM sodium citrate (J.T. Baker cat# 3646-01) at 30-35°C for 15 minutes, and then were cooled for another 15 minutes, followed by a 1 hour 50mM glycine (Sigma, cat# G-7126, in PBS) incubation. We then performed a blocking step in a solution of 5% bovine serum albumin (BSA, Jackson ImmunoResearch, cat# 001-000-162), 10% normal goat serum (NGS, Jackson ImmunoResearch cat# 005-000-121), 0.4% Triton X-100, 0.1% acetylated bovine serum albumin (BSA-c, Aurion, Electron Microscopy Sciences, cat#25557) in 0.1M phosphate buffer (PB) for 1 hour. Sections were then incubated for 72 hours at 4°C in the rabbit primary antibody for NMDAR-GluN2B at 1:100 in antibody dilution buffer, which was composed of 1% BSA; 1% NGS; 0.1% BSA-c; and 0.1% Aurion coldwater fish gelatin (CWFG Electron Microscopy Sciences cat# 25560) in PB. Sections were then incubated in Aurion F(ab) fragment of goat anti-rabbit ultrasmall at 1:50 (Electron Microscopy Sciences, cat#25361) at 4°C overnight. Then, the sections underwent a 4% paraformaldehyde (in PBS) postfix for 5 minutes, followed by a 10 minute 50mM glycine incubation, and then a few quick distilled water washes. Silver enhancement was performed with the Nanoprobes HQ Silver kit (cat# 2012-45ML) in the dark for 20-30 minutes, and produced variable sized particles. A control section was run in tandem, with the sole difference being omission of the primary antibody from the antibody dilution buffer. When imaged, gold particles were extremely rare, even at the edge of the tissue where labeling is typically dense and noisy, indicating that the secondary antibody was highly specific for the primary antibody (<u>Fig. S6C-D</u>).

#### Double label immunoEM for NMDAR-GluN2B and MAP2

To perform double-labeling for NMDAR-GluN2B and MAP2, we paired NMDAR-GluN2B immunogold with immunoperoxidase non-nickel diaminobenzidine (DAB) for MAP2. We performed the antigen retrieval and glycine incubation as above. Then, sections underwent a 30 minute 0.3% hydrogen peroxide incubation at 4°C and avidin-biotin blocking (Vector cat# SP-2001) to prevent non-specific labeling by the immunoperoxidase product. Then sections were preblocked as above, and incubated in the primary antibody for MAP2 for 48 hours at 4°C. They were then washed and incubated with biotinylated goat anti-chicken (Jackson ImmunoResearch, cat# 103-065-155) at 1:200 for 3 hours at room temperature. Sections then underwent incubation with avidin biotin complex (ABC, Vector cat# PK-6100) for 1.5 hours, and visualized using a DAB kit (Vector cat# SK-4100, nickel excluded). Following that, we performed washes and followed the procedures for single-label immunogold labeling as above for the NMDAR-GluN2B-gold. Initially, we tried reversing the order of operations, with the immunogold first, followed by the immunoperoxidase, and found that the gold particles fared better when performed as the second step.

#### EM Processing

After immunolabeling, sections were processed for electron microscopy. They were post-fixed in 4% paraformaldehyde in 0.1M PBS for 20 minutes, and then submersed in 1% osmium tetroxide in PB, which was then immediately diluted to 0.5%, and the sections incubated in the dark for 30 minutes at 4°C. Then, following PB washes, sections were washed in 3 x 5-10 minute sets of washes in ascending 50% and 70% ethanol dehydration steps, before being incubated in 1% uranyl acetate in 70% ethanol for 40 minutes. Sections were further dehydrated in 3×5 minute 95% and 100% ethanols, followed by propylene oxide washes (Electron Microscopy Sciences cat#20401). Finally, sections were infiltrated with Durcupan resin (Electron Microscopy Sciences cat#14040), and baked at 60°C for 72 hours sandwiched between sheets of Aclar (Electron Microscopy Sciences, cat# 50425).

### Immunohistochemical procedures for multi-label immunofluorescence

To label NMDAR-GluN2B and the CBPs PV, CB, and CR, we performed antigen retrieval on free-floating sections, as above, except at 75-80°C, and doubled the time for the water bath and cooling steps. We then performed the glycine and blocking steps, as above. The antibody dilution buffer was the same as above, but excluding the CWFG and substituting normal donkey serum (Jackson ImmunoResearch, cat# 017-000-121). Primary incubation occurred in antibody dilution buffer for NMDAR-GluN2B, PV, CB, and CR for 72 hours at 4°C. Then, we incubated the sections in species-specific AlexaFluor donkey secondary antibodies at 1:100 (Invitrogen, AlexaFluor-568 anti-mouse cat# A10037, AlexaFluor-488 anti-rabbit cat# A32790, AlexaFluor405 anti-goat cat# A48259) and a biotinylated donkey anti-guinea pig (Jackson ImmunoResearch cat# 705-065-148) for 3 hours at 4°C in the dark. Then, sections were washed and incubated in Streptavidin-647 (Invitrogen, cat# S21374) at 1:200 for 3 hours at room temperature. Sections were mounted using ProLong Gold Antifade Mountant (Invitrogen, cat# P36930). We also ran a preadsorption control test for NMDAR-GluN2B. One section was run in tandem with the above, but incubated in antibody dilution buffer containing the Alomone NMDAR-GluN2B primary antibody, but with the addition of the blocking peptide for the NMDAR-GluN2B antigen (Alomone, cat# BLP-GC003), at 10x the concentration of the primary antibody (<u>Fig. S6A-B</u>). The sections were incubated in a goat anti-rabbit Alexafluor secondary antibody (Invitrogen, cat# A11008).

### Imaging procedures, 2D Sampling, and Analysis for immunoEM

#### Imaging and Sampling

Two or more sections were processed per animal per area for single-labeled NMDAR-GluN2B analysis, and for the double-labeling MAP2/NMDAR-GluN2B analysis. For the single-label NMDAR-GluN2B analysis, a minimum 4 blocks, 2 from each section, were examined per area per subject for sampling. One block per section per area were analyzed for the double-labeled MAP2/NMDAR-GluN2B analysis. Blocks were dissected from layer III within the principal sulcus of the dlPFC, and along the medial wall of the SGC, and mounted on Durcupan resin blocks, and then sectioned at 50nm on an ultramicrotome (Leica). Short series (10-20 sections, typically) were collected on Butvar-coated (Electron Microscopy Sciences, cat# 11860) copper slot grids near the top of the section. Grids were imaged using a Talos L120C transmission electron microscope (Thermo Fisher Scientific). We mapped each section and outlined the boundaries of the antibody penetration region, which is typically reliable within 5-40 nm from the edge of the tissue section, but can be variable by section depending on plane of cut. Using a meander-scan approach, we used a systematic sampling acquisition protocol, snapping every 2^nd^ or 3^rd^ field of view (depending on extent of antibody penetration zone) as we traversed the antibody penetration zone of the section, at 11-13,000x magnification using a Ceta CMOS camera. For single-labeled NMDAR-GluN2B immunogold sections, approximately 300-475 images were sampled for per cortical area per subject, depending on immunolabeling density (Monkey 1: A25, 293 images, dlPFC, 475 images; Monkey 2: A25, 305 images, dlPFC 346 images). For double-labeled MAP2/NMDAR-GluN2B sections, approximately 200-500 images were acquired for analysis per area per case, depending on immunolabeling density (Monkey 1: A25, 203 images, dlPFC, 289 images; Monkey 2: A25, 550 images, dlPFC, 292 images). Images were adjusted for brightness and contrast using Adobe Photoshop CS5 Extended (version 12.0.4 ×64, Adobe Systems Incorporated) for figures.

#### Analysis of 2D sampled electron micrographs

All 2D images were analyzed using Reconstruct (Fiala, 2005). We examined all gold particles in all images, classified each particle’s parent structure as a spine, axon, bouton, dendrite, likely glial process, cell soma or undetermined structure using classical criteria (Peters et al., 1991). Likely because of differences in fixation conditions between the subjects, the total proportion of undetermined structures was variable across subjects, in particular for small or broken structures, or for thin structures with mitochondria that did not have synaptic interactions, spines, or telltale indicators of axonal, glial, or dendritic identity. We classified immunogold particles as cytoplasmic if not touching any external membranes, and synaptic if found touching the post-synaptic density (PSD) of a synapse. We classified immunogold particles as perisynaptic if the immunogold particle was within approximately <100 nm from the synapse along the membrane, and extrasynaptic if the immunogold particle was >100 nm from the PSD along the membrane (Hardingham and Bading, 2010; Petralia et al., 2010) though the extent of the perisynaptic region appears to be somewhat loosely defined (Newpher and Ehlers, 2008; Gladding and Raymond, 2011). The total number of Glun2B+ spines sampled varied by area by case, depending on immunolabeling success and plane of sectioning (Monkey 1: A25, 229 spines, dlPFC 483 spines; Monkey 2: A25, 242 spines, dlPFC 162 spines). After tracing the outline of each spine head containing positive NMDAR-GluN2B immunolabeling, we computed the major Feret’s diameter of each spine to determine if there were detectable differences in NMDAR-GluN2B+ spine head size across cortical areas. To determine if we could detect differences among randomly sampled NMDAR-GluN2B-negative spine heads across areas, we selected one image sampling session from each cortical area per subject, and measured all spines present that were not NMDAR-GluN2B+, which we call “neuropil” spines for comparison to NMDAR-GluN2B+ spines. Given that we performed only a 2D analysis, it is important to note that these spines may have had NMDAR-GluN2B immunogold particles in planes outside the imaged section, and thus the type II error rate (false negative) is elevated for this analysis, though it represents a systematic bias equivalently present across the compared cortical areas (NMDAR-GluN2B-negative spines sampled, Monkey 1: A25, 113 spines, dlPFC 153 spines; Monkey 2: A25, 310 spines, dlPFC 124 spines). For all membrane-bound immunogold particles in spines, we measured the shortest distance from the membrane to the smooth endoplasmic reticulum (SER) in plane, making multiple measurements from the site of gold contact with the membrane and the closest elements of the SER in plane. Given that there may be elements of the SER outside the sectioning plane that may have been actually closer to the immunogold particles (e.g., just above or below), this analysis may also contain an elevated type II error rate.

### Imaging procedures, Sampling, and Analysis for MLIF

Sections were imaged on a Zeiss LSM 880 Airyscan with the Plan-Apochromat 20x/0.8 M27 objective. Z-stacks were obtained with ∼1-µm steps through the depth of the tissue under laser excitation at 405nm, 488nm, 561nm, or 633nm. Emission filter bandwidths and sequential scanning acquisition were set up in order to avoid spectral overlap between fluorophores. Confocal images were deconvolved with Huygens Professional version 22.04 (Scientific Volume Imaging, The Netherlands) using uniform parameters. For quantitative analysis in the dlPFC, we located the principal sulcus, and used the NMDAR-GluN2B labeling in pyramidal neurons, and the distribution of inhibitory neurons, which differentially populate laminar compartments (Joyce et al., 2020; Medalla et al., 2023) to locate layer III. We then systematically sampled from deep layer III in parallel to layer I, acquiring an image in every other field of view in dlPFC along the axis of the principal sulcus; we sampled exhaustively in the medial wall of A25, given its shorter breadth. Two sections were sampled per cortical area per case. From each stack we extracted maximum projections of a few sections near the top and bottom of the stack for quantitative analysis (Monkey 1: A25, 12 images; dlPFC, 17 images; Monkey 2, A25, 17 images; dlPFC, 15 images). We analyzed these maximum projections using QuPath 0.5.0 (Bankhead et al., 2017)). If the images did not have uniform illumination, we analyzed the image in pieces containing uniform illumination. We first manually segmented the PV, CB, and CR inhibitory neurons, and CB+ pyramidal neurons (Kondo et al., 1999; Joyce et al., 2020; Datta et al., 2024). In the isolated NMDAR-GluN2B channel, we traced any pyramidal neurons that were negative for CB+, and sampled from immunonegative “neuropil” segments of tissue containing no obviously labeled processes, for background calibration. We measured the mean intensity (MI) using 0.1 µm pixel size. We averaged the MI across immunonegative background regions, and across the all NMDAR-GluN2B+ pyramidal neurons. We then used these numbers to create a ceiling and floor for binning all inhibitory neurons. We deemed a CBP+ neuron NMDAR-GluN2B negative if it had an MI at or below the average across sampled background regions, and NMDAR-GluN2B strong if it had an MI at or above the mean across NMDAR-GluN2B+ pyramidal neurons (including pyramidal neurons also positive for CB), and included a few intermediate bins. We then averaged the proportion falling into each bin across cases. See <u>Figure S5</u> for these data. Total numbers of neurons were variable across areas, (Monkey 1: A25, 383 CBP+ inhibitory neurons and 719 CB/NMDAR-GluN2B+ pyramidal neurons, dlPFC, 452 CBP+ inhibitory neurons, 254 CB/NMDAR-GluN2B+ pyramidal neurons; Monkey 2: A25, 864 CBP+ inhibitory neurons and 834 CB/NMDAR-GluN2B+ pyramidal neurons, dlPFC, 449 CBP+ inhibitory neurons, 97 CB/NMDAR-GluN2B+ pyramidal neurons) given that these cortical areas have variable densities of these neurons (Joyce et al., 2020). Images were adjusted for brightness and contrast using Adobe Photoshop CS5 Extended (version 12.0.4 ×64, Adobe Systems Incorporated) for figures.

### Statistical Analysis

Statistical analysis and plot preparation for figures was performed in Prism (Graphpad). We assessed the properties of our distributions and determined the appropriate parametric or nonparametric tests. To analyze the membrane distribution of NMDAR-GluN2B particles, average spine sizes, and distances of immunogold particles to the SER, we averaged across Monkey 1 and Monkey 2 and performed a one-way ANOVA with post-hoc Tukey tests. Figures were prepared in Adobe Illustrator 25.0.1 (Adobe Systems Incorporated version 27.7).

## Results

### NMDAR-GluN2B is more frequently synaptic in dlPFC spines, and extrasynaptic in SGC spines

To study the membrane localization of NMDAR-GluN2B in SGC and dlPFC, we used pre-embedding immunoEM to label NMDAR-GluN2B with immunogold particles, and performed systematic 2D sampling in layer III of the SGC and dlPFC. Spines found in layer III of both areas were frequently NMDAR-GluN2B+ using both post-embedding and pre-embedding techniques (<u>Figs. 1-2</u>). In our pre-embedding preparation (<u>Figs. 2, S1</u>) we most often found NMDAR-GluN2B immunolabeled particles in the cytoplasm, which we have interpreted as intracellular trafficking events. Synaptic NMDAR-GluN2B were readily observable in SGC and dlPFC spines (<u>Figs. 2A1,B1-B2; S1A,E</u>). We also observed NMDAR-GluN2B in membranes outside the synapse at extrasynaptic and perisynaptic locations (<u>Figs. 2A2-A3,B3; S1B-D,F-H</u>). We pooled the data from all spines within each cortical area per monkey, and quantified the proportion of NMDAR-GluN2B immunogold particles found at synaptic, perisynaptic, extrasynaptic and cytoplasmic locations (<u>Fig. 2C</u>). <u>Figure 2C</u> demonstrates that both monkeys had a higher proportion of extrasynaptic NMDAR-GluN2B in SGC spines, while NMDAR-GluN2B immunogold particles were more likely to be found in the synapse of dlPFC spines. <u>Figure 2D</u> shows that when averaged, these relationships were significant, indicating that among membrane-bound immunoparticles, the ratio of extrasynaptic to synaptic NMDAR-GluN2B particles was just over 2:1 in A25 spines, while it was 0.9:1 in dlPFC spines.

**Figure 2.**
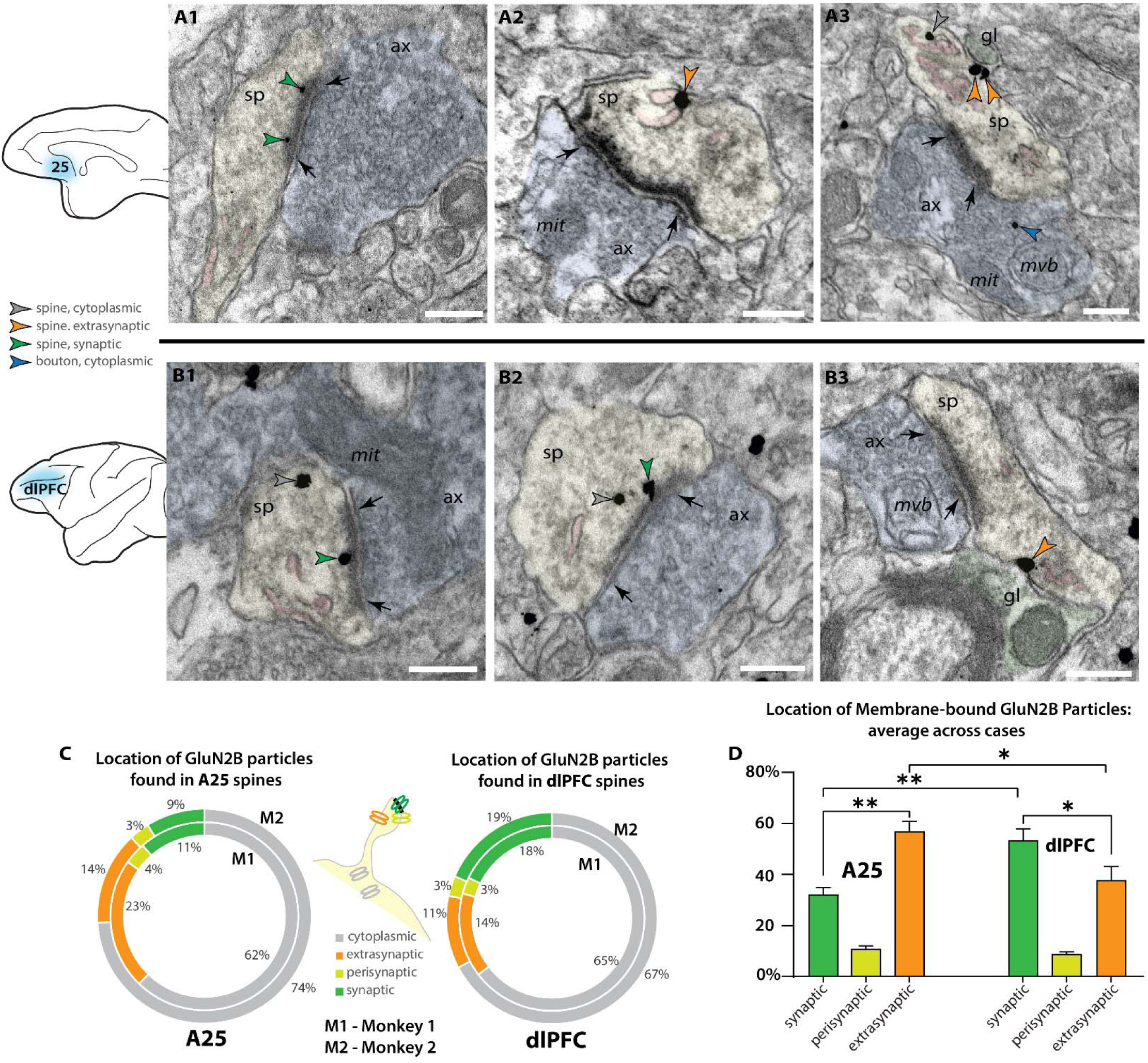
Higher proportion of extrasynaptic GluN2B gold particles in layer III A25 spines than dlPFC spines. ***A***, Electron micrograph depicting spines in A25 layer III. ***A1***, An A25 spine (sp, pseudocolored yellow) with NMDAR-GluN2B immunogold particles (green arrowheads) in the postsynaptic density of a synapse (black arrows) formed by an axonal bouton (ax, pseudocolored blue). The spine contains a spine apparatus, an extension of the smooth endoplasmic reticulum (SER, pseudocolored pink) in the spine neck. ***A2-A3,*** A25 spines with NMDAR-GluN2B adhered to extrasynaptic membranes (orange arrowheads), near the SER. In A3, the extrasynaptic NMDAR-GluN2B is apposed to a structure consistent with glial morphology (gl, pseudocolored green), and the bouton contains a presynaptic cytoplasmic NMDAR-GluN2B (blue arrowhead) among the vesicles. ***B,*** Electron micrographs depicting spines in dlPFC layer III. ***B1-B2,*** dlPFC spines containing synaptic NMDAR-GluN2B (green arrowheads), and cytosolic NMDAR-GluN2B (grey arrowheads), which are likely being trafficked. ***B3,*** A dlPFC spine with an extrasynaptic NMDAR-GluN2B in the spine neck, apposed to a structure consistent with glial morphology, and near a spine apparatus in the spine neck. ***C,*** Nested pie charts depicting location of NMDAR-GluN2B immunogold particles in spines of A25 (left), and dlPFC (right) in Monkey 1 (M1, inside) and Monkey 2 (M2, outside). Percent of NMDAR-GluN2B immunogold particles found in the cytoplasm (grey), post-synaptic density (green), perisynaptic membrane (yellow-green), and extrasynaptic membrane (orange) of GluN2B+ spines. ***D,*** Plot depicting the location of membrane-bound GluN2B immunogold particles, in relation to the synapse, averaged across M1 and M2. Error bars depict standard deviation. One-way ANOVA, F(5,6=67.71, p<0.001, with post-hoc Tukey test). *, p < 0.05; **, p< 0.01, *** p< 0.001; ****, p<0.0001; scale bars, 200nm. ax,axon; gl, glial process; mit, mitochondria; mvb, multivesicular body; SER, smooth endoplasmic reticulum spine apparatus; sp, spine.

We also performed supplemental analyses to characterize the spines that expressed NMDAR-GluN2B. We measured the major Feret’s diameter of each GluN2B+ spine, and in a smaller set of sampled images per subject and cortical area we also measured all other GluN2B-spines present in the images (<u>Fig, S2A</u>). The mean major diameter of GluN2B+ spines across the two cortical areas was similar, approximately ∼0.6 µm. The mean diameter of GluN2B+ spines was larger for both animals than were GluN2B-spines, with the caveat that some GluN2B-spines may contain GluN2B in planes above or below the plane sampled in our analysis. We also measured the distance of each membrane-bound NMDAR-GluN2B immunogold particle to any evident SER spine apparatus in plane. Many NMDAR-GluN2B were within tens of nanometers from the SER, indicating that they are within physiological range to evoke calcium-mediated calcium release from the SER (<u>Fig. S2B</u>), as demonstrated in other systems (see (Datta et al., 2024)). Given the constraints of our 2D sampling paradigm, many of these membrane-bound NMDAR-GluN2B immunogold particles may be closer to SER elements that were present in unsampled portions of the spine above and below the plane of section. We also observed presynaptic NMDAR-GluN2B labeling (<u>Fig. S3</u>), although these gold particles were very rarely adhered to the membrane (<5% in boutons sampled in both dlPFC and A25). They were mostly found in the cytoplasm amidst the vesicles.

### NMDAR-GluN2B is more likely to be extrasynaptic in A25 than dlPFC putative excitatory dendrites

In addition to spines, we also characterized NMDAR-GluN2B expression in dendritic shafts that were contained in our sampled images. We were able to identify some dendrites that had spines in plane, indicating they were likely the dendrites of excitatory neurons (<u>Fig. 3A2-3,B1</u>), given that cortical excitatory neurons are spiny (Peters et al., 1991; Hsu et al., 2017). To supplement this analysis, we then turned to double label immunoEM to evaluate excitatory dendritic shafts, using NMDAR-GluN2B and MAP2, a protein prominently expressed in the dendrites of pyramidal neurons in monkey PFC (e.g., (Tsolias and Medalla, 2021)), although we have recently observed that MAP2 can be expressed in proximal dendrites of inhibitory neurons in dlPFC (Joyce et al., 2024a). We then performed systematic sampling in tissue double-labeled for MAP2 and GluN2B. In MAP2+ dendrites, synapses formed on dendritic shafts were very rare, and in only one instance we observed a MAP2+ dendrite containing more than one synapse on the dendritic shaft in plane, indicating that it may have been an inhibitory dendrite. Given that the majority of asymmetric presumed excitatory synapses formed on prefrontal pyramidal neurons occur on spines rather than dendritic shafts (Hsu et al., 2017), the rarity of dendritic shaft synapses found in our sampled MAP2+ dendrites suggests that the majority of them were likely excitatory dendrites. Among MAP2+ dendrites (*e.g.*, <u>Fig. 3A1,B2, S4</u>), we counted the gold particles and quantified their location (<u>Fig. 3C, S4</u>). In A25, NMDAR-GluN2B were more frequently found in the membranes of dendritic shafts than in dlPFC, and this relationship was significant (<u>Figs. 3D, S4</u>).

**Figure 3.**
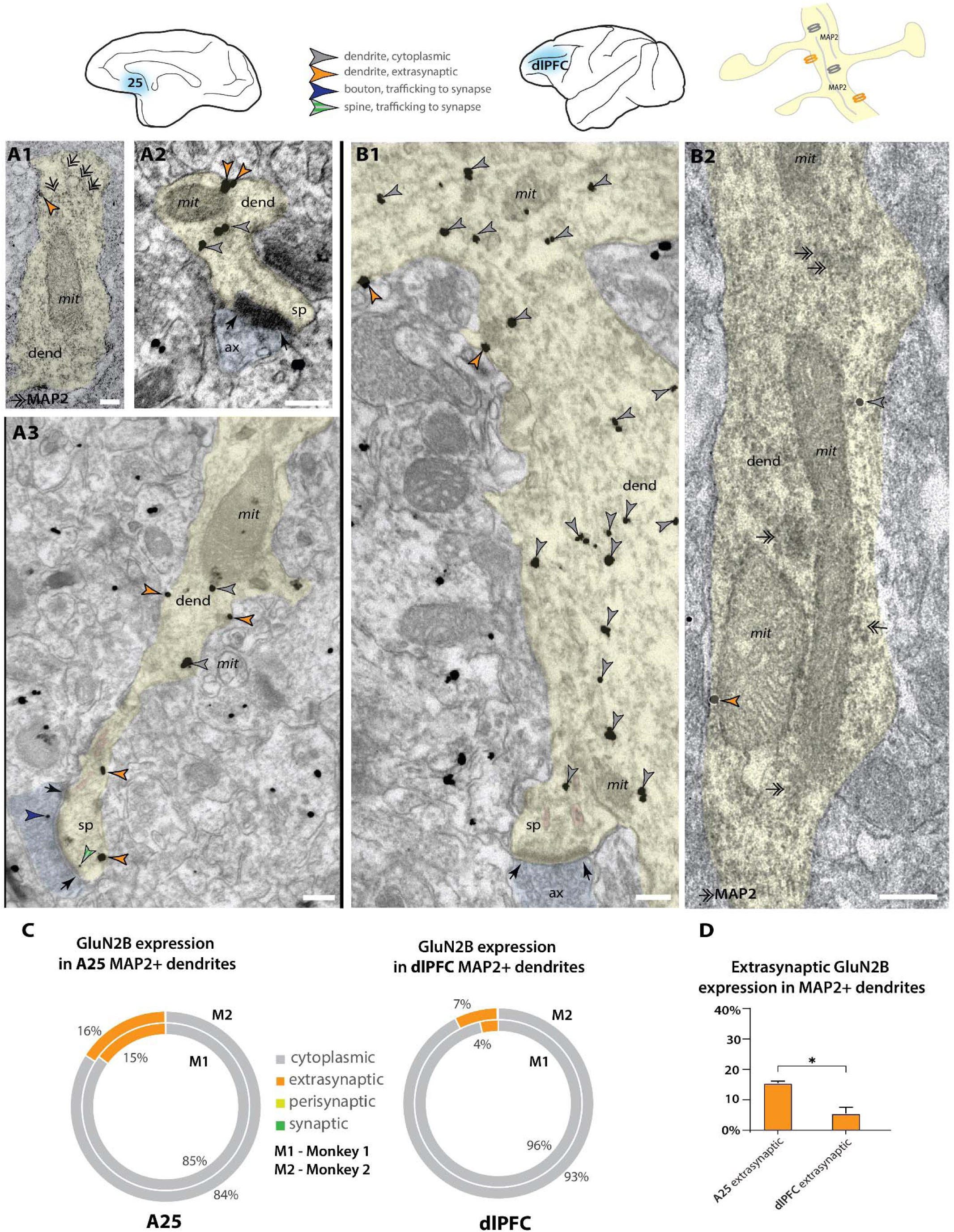
Higher extrasynaptic expression in dendrites of putative excitatory neurons of A25 compared to dlPFC. ***A,*** Electron micrographs from layer III A25. ***A1***, A putative excitatory dendrite, labeled with MAP2+ non-nickel immunoperoxidase diaminobenzidine (smudge-like precipitate, double-headed arrows), expressing an extrasynaptic NMDAR-GluN2B (orange arrowhead); ***A2-A3***, NMDAR-GluN2B at extrasynaptic (orange arrowheads), cytoplasmic (grey arrowheads), or near-synaptic locations (grey-green striped arrowhead) in putative excitatory dendrites. ***B***, Electron micrographs from layer III dlPFC. ***B1,*** A putative excitatory dendrite, with a spine in plane, expressing cytoplasmic and extrasynaptic NMDAR-GluN2B. ***B2,*** A putative excitatory dendrite, labeled with MAP2, expressing extrasynaptic and cytoplasmic NMDAR-GluN2B. ***C,*** Nested pie charts depicting the percent of NMDAR-GluN2B immunogold particles found at cytoplasmic, extrasynaptic, perisynaptic, and synaptic locations in MAP2+ dendritic shafts in A25 (left) and dlPFC (right) of Monkey 1 (outside) and Monkey 2 (inside). Synapses on the shaft of MAP2+ dendrites were rare, and synaptic NMDAR-GluN2B on MAP2+ shaft synapses were extremely rare (<0.2% of all immunogold particles in all areas analyzed). ***D,*** Mean percent of extrasynaptic NMDAR-GluN2B immunogold particles across all MAP2+ dendrites in A25 and dlPFC. One-way ANOVA with post-hoc Tukey test, F(3,4) = 1691, p<0.0001. *, p<0.05; pink pseudocolor, SER spine apparatus; black arrows, synapse; scale bars, 200nm; ax, axon; dend, dendrite; MAP2, microtubule-associated protein-2; mit, mitochondria; sp, spine;

### Prominent somatic extrasynaptic NMDAR-GluN2B expression on A25 pyramidal-like neurons

Membrane expression of NMDAR-GluN2B at the soma occurred in both A25 and the dlPFC, though this was difficult to quantify because cell identity was not always determinable in our 2D sampled images. In one A25 section, we found a pyramidal-like somata with an apical dendrite, basal dendrite, and axon initial segment well contained within the antibody penetration zone. The pyramidal-like neuron had robust NMDAR-GluN2B labeling in the cytoplasm of the soma and proximal dendrites and at extrasynaptic sites (<u>Fig. 4</u>), in contrast to sparser NMDAR-GluN2B labeling in surrounding neuropil (<u>Fig. 4H</u>). Notably, we observed robust NMDAR-GluN2B in the nucleus and in the nuclear membrane (<u>Fig. 4H</u>).

**Figure 4.**
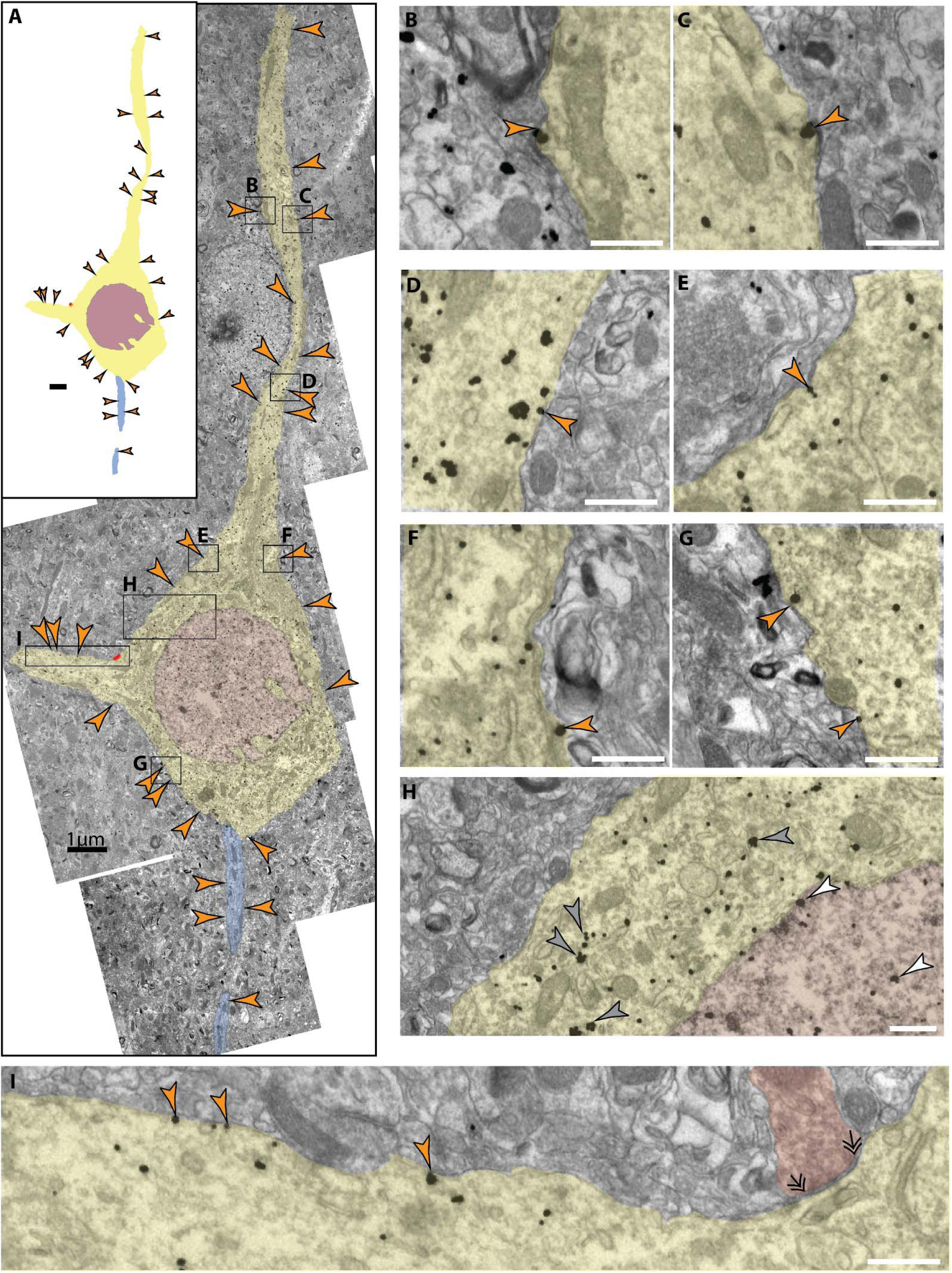
Extrasynaptic NMDAR-GluN2B expressed at a pyramidal-like soma in layer III A25. ***A,*** A panorama of stitched electron micrographs depicting a pyramidal-like soma and dendrites (psuedocolored yellow), including nucleus (pseudocolored plum) and its axon initial segment (pseudocolored blue). NMDAR-GluN2B are prevalently expressed in extrasynaptic membranes (orange arrowheads) at the soma and proximal processes. Black boxes depict inset locations for subsequent panels. ***B-G,*** Insets from **A** depicting extrasynaptic NMDAR-GluN2B at higher magnification. ***H,*** Inset from **A** selected to emphasize the antibody labeling specificity of the labeled neuron compared to the surrounding neuropil, as well as to emphasize nuclear labeling. Few NMDAR-GluN2B immunoparticles are evident in the surrounding neuropil, while the pyramidal-like soma densely expresses cytosolic NMDAR-GluN2B immunoparticles (grey arrowheads). NMDAR-GluN2B immunoparticles are also present in the nucleus (pseudocolored plum, white arrowhead). ***I,*** Inset from A depicting several more extrasynaptic NMDAR-GluN2B on a basal dendrite, as well as a symmetric synapse (double arrowheads) formed on the soma (axon pseudocolored red). Scale bars in **B-I**, 200nm.

### High likelihood inhibitory dendrites have less synaptic NMDAR-GluN2B expression than spines, as well as extrasynaptic expression

We also examined the expression of NMDAR-GluN2B in putative inhibitory dendrites. Inhibitory dendrites in the cortex are predominantly aspiny or sparsely spiny, and receive most synaptic input on the dendritic shaft (Peters et al., 1991). We classified inhibitory dendrites as “high likelihood” based on the presence of two or more shaft synapses in plane, and no spines in plane. <u>Figure 5A-B</u> depicts examples of these dendrites in A25 and dlPFC. NMDAR-GluN2B found in the postsynaptic density (<u>Fig. 5C</u>, 1-3% across areas) occurred at a much lower frequency than those observed in spines found in the same sampled images (∼10% synaptic NMDAR-GluN2B in A25 spines vs. ∼20% in dlPFC, <u>Fig. 2C</u>). In both areas, the extrasynaptic proportion was slightly above 10%, and there were no differences detected across areas when the subjects were pooled. When NMDAR-GluN2B was found in the PSD, the immunogold particles were most often affiliated with the far edge of the PSD and rarely in the main body of the synapse (e.g., <u>Fig. 5A3, B2</u>).

**Figure 5.**
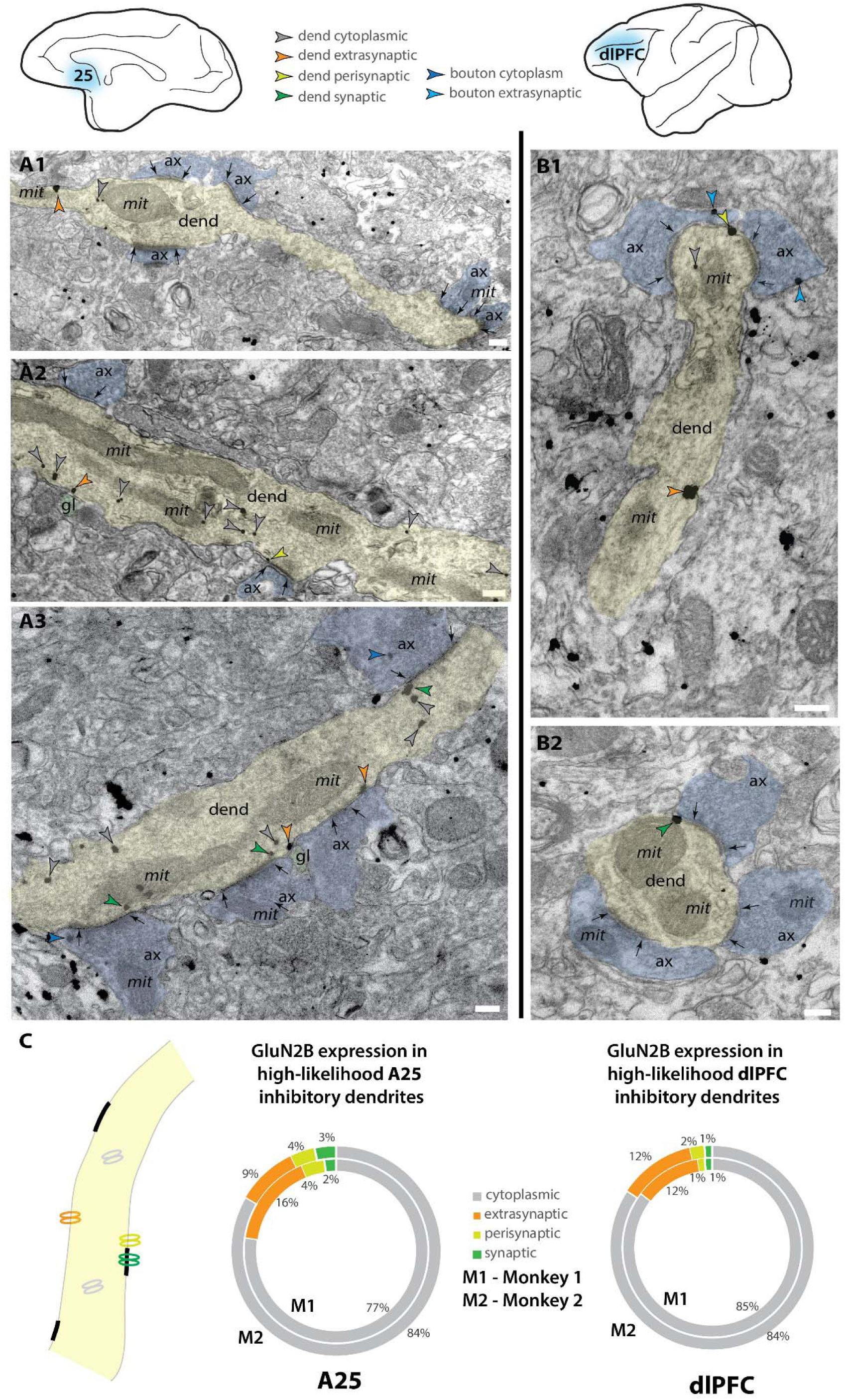
NMDAR-GluN2B are expressed in inhibitory dendrites in layer III A25 and dlPFC. Electron micrographs depicting high likelihood inhibitory dendrites in A25 (**A**) and dlPFC (**B**). Dendrites were deemed high likelihood inhibitory dendrites by the lack of spines in plane, and the presence of two or more asymmetric synapses formed on the dendritic shaft, as inhibitory dendrites in the cortex are sparsely spiny or aspiny (Peters et al., 1991). ***A1-A3,*** Examples of NMDAR-GluN2B immunogold labeling in high likelihood inhibitory dendrites found in layer III A25. ***B1-B2,*** Examples of NMDAR-GluN2B immunogold labeling in high likelihood inhibitory dendrites found in layer III dlPFC. ***C,*** Nested pie charts depicting location of NMDAR-GluN2B immunogold particles in high likelihood dendrites of A25 (left) and dlPFC (right) for Monkey 1 (inside) and Monkey 2 (outside). No statistically significant differences were detected between dlPFC and A25. Scale bars, 200nm. ax, axon; dend, dendrite; gl, glial process; mit, mitochondria.

### NMDAR-GluN2B expression is roughly equivalent across inhibitory neuron somata in both areas

We were also interested in whether the NMDAR-GluN2B expression level varied across inhibitory neuron types. In primates, the calcium-binding proteins (CBP) are a useful way to classify inhibitory neurons. The CBPs parvalbumin (PV), calbindin (CB), and calretinin (CR) label upwards of 85% of all cortical inhibitory neurons and are non-overlapping, that is they are neurochemically distinct (Conde et al., 1994; DeFelipe, 1997; Medalla et al., 2023). To measure whether there were expression level differences across CBPs, we used multi-label immunofluorescence (MLIF) for PV, CB, CR, and NMDAR-GluN2B (<u>Figs. 6A-B, S5A-B</u>). We segmented traces for the somata of the inhibitory neurons, and then in the isolated GluN2B channel, we traced the outlines of pyramidal-like neurons, and sampled immunonegative “background” regions of the tissue that contained no labeled processes. We measured the mean intensity (MI) for each trace, and then compared the inhibitory neuron traces to the MI found in pyramidal neurons and in “background”-like immunonegative areas (<u>Fig. S5C-F</u>). This allowed us to create expression bins per sampling site, and then combine information on the bins across sampling sites (<u>Fig. S5G</u>). <u>Figure 6C-D</u> depict the results of this analysis. The large majority of inhibitory neurons (∼85%) in both areas expressed NMDAR-GluN2B at an intensity level below their nearby pyramidal neurons, and 20% or less of each inhibitory neuron type were negative for NMDAR-GluN2B. There were no detectable differences across areas or CBP types.

**Figure 6.**
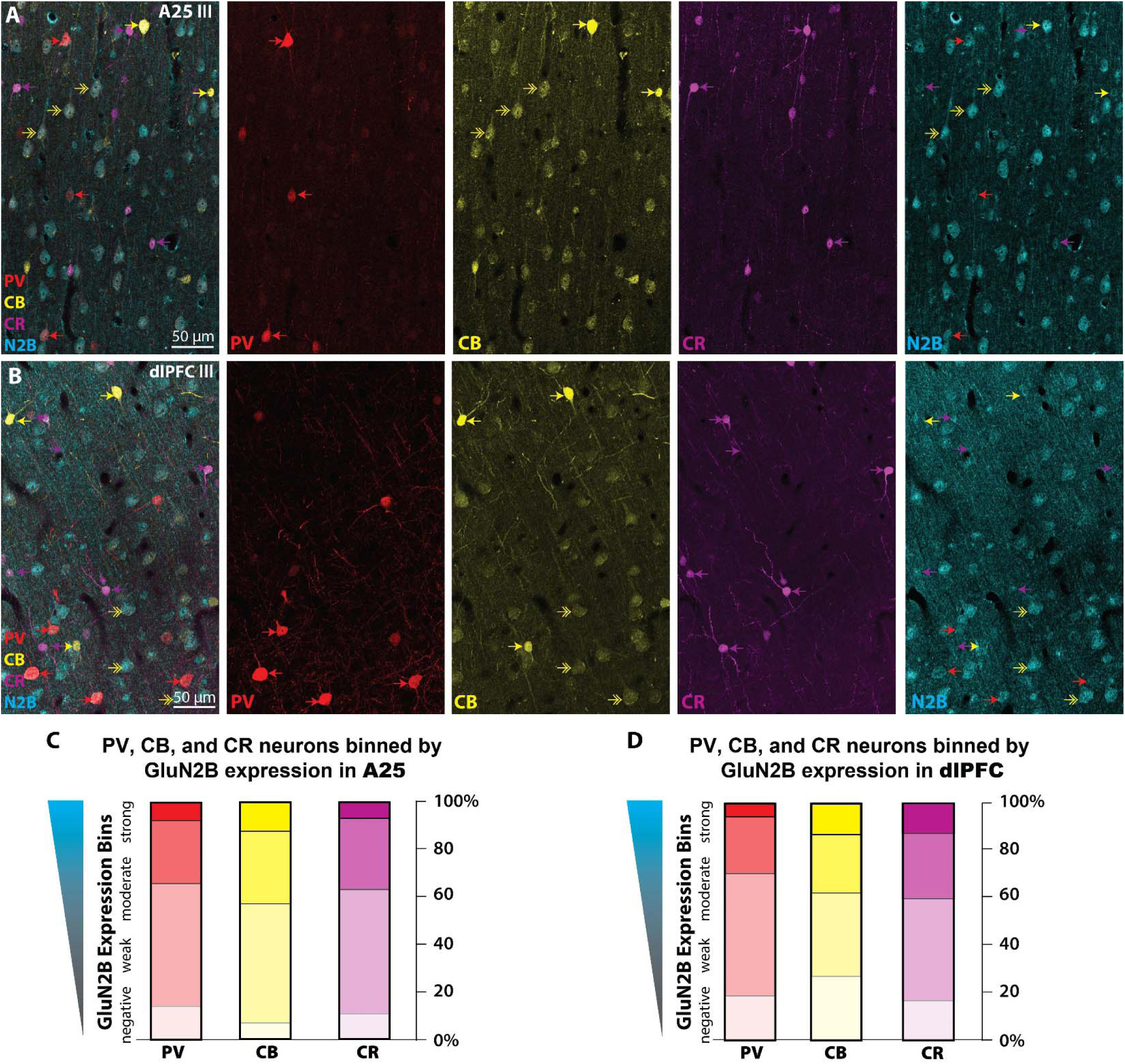
NMDAR-GluN2B are expressed at equivalent levels across CBP+ inhibitory neurons in A25 and dlPFC. ***A-B***, Maximum projection Images of layer III A25 (**A**) and dlPFC (**B**) obtained via confocal microscopy depicting multiple immunofluorescence labeling for PV (red), CB (yellow), CR (magenta), and NMDAR-GluN2B (cyan). Color-coded arrows depict inhibitory neurons, and double-headed arrows indicate CB+ pyramidal neurons. ***C-D,*** Stacked bar charts depicting mean proportion of PV (red), CB (yellow), and CR (magenta) that fell into negative, weak, moderate or strong NMDAR-GluN2B expression bins as determined by the mean intensity (MI) in the NMDAR-GluN2B channel. GluN2B-negative is defined by MI at or below the average amount in immunonegative sampled neuropil regions. GluN2B-strong is defined by MI at or above the average found in morphologically identified pyramidal-like neurons expressing NMDAR-GluN2B. See Supplemental Figure 6 for more detailed information about the inhibitory neuron analysis. CB, calbindin; CR, calretinin; PV, parvalbumin.

## Discussion

The current study found variation in the synaptic vs. extrasynaptic expression of NMDAR-GluN2B in dendrites and spines of layer III of the macaque SGC and dlPFC. Our data suggest that there is a higher proportion of NMDAR-GluN2B expressed at extrasynaptic membrane locations among putative pyramidal neurons in SGC than dlPFC, while synaptic NMDAR-GluN2B predominated in dlPFC. As the SGC appears to be relatively enriched in NMDAR-GluN2B (Burt et al., 2018) --although this pattern may be less pronounced in rhesus macaques (Chen et al., 2023)-- our data suggest that there is a substantial population of extrasynaptic NMDAR-GluN2B in SGC compared to some other regions of the PFC. The preponderance of synaptic NMDAR-GluN2B in dlPFC is consistent with previous work demonstrating synaptic GluN2B in this region, and demonstrating the importance of NMDAR-GluN2B to dlPFC neuronal firing during working memory (Wang et al., 2013a). This contrasts with putative inhibitory neurons in both areas, where NMDAR-GluN2B membrane expression was rarely synaptic, with no detectable areal differences among inhibitory-like dendrites sampled in this study. When present, synaptic NMDAR-GluN2B in inhibitory neuron dendrites were found at the periphery of the PSD.

Our study utilized pre-embedding immunoEM, which entails the binding of antibodies to antigens in free-floating sections that are tens of microns in thickness, followed by processing for EM, embedding in resin, and ultramicrotomy to section into ultrathin sections for EM imaging. This is in contrast to post-embedding immunoEM, in which immunohistochemical procedures are performed on sections tens of nanometers in thickness, from blocks that have already been processed for EM and embedded in resin (e.g., <u>Fig. 1</u>). We use pre-embedding immunoEM for examining molecular localization in organelles and extracellular membranes because of its gentler treatment of tissue ultrastructure, allowing the ability to access extrasynaptic compartments more readily given their preservation. However, pre-embedding immunoEM has its caveats, such as limited antibody penetration, and the difficulty of achieving full antibody access to the dense matrix of proteins in the PSD (reviewed in (Petralia and Wang, 2021)). For this reason, pre-embedding immunoEM likely undersamples from the PSD, a critical constraint for our study, which means that our data should not be interpreted as ground truth absolute expression data for synaptic vs. extrasynaptic expression, but rather a dataset that can reveal comparative relationships between cortical areas subjected to the same labeling and analytical procedures. This methodological difference likely explains in part the higher NMDAR-GluN2B synaptic presence that was observed in an earlier study of dlPFC (Wang et al., 2013a), which may be compounded by the possibility of differences in antibody performance. The critical piece of evidence that we offer is that it appears that the detectable membrane expression patterns can shift across primate PFC areas under equivalent labeling procedures, with increasing extrasynaptic expression in the SGC as compared to the dlPFC.

Extrasynaptic NMDAR likely serve important functions in normal homeostatic physiological conditions. For example, studies in rodent hippocampal cultures and *ex vivo* slice preparations suggest that extrasynaptic NMDAR are stored for shuttling and trafficking in and out of synapses, even for those NMDAR that are outside the loosely defined perisynaptic region [(Kortus et al., 2023); reviewed in (Groc and Choquet, 2020; Petit-Pedrol and Groc, 2021)]. Extrasynaptic NMDARs can propagate dendritic spikes after an initial depolarizing event according to early theoretical modeling (Rhodes, 2006) and more recent slice electrophysiology studies in rodents, e.g., the mPFC (Chalifoux and Carter, 2011), and human medial temporal cortex (Testa-Silva et al., 2022). Dendritic NMDA spikes play a role in integrative processes and signal amplification *in vivo* in mouse somatosensory cortex (Palmer et al., 2014). Extrasynaptic NMDAR have also been linked with astrocytic glutamate release, producing slow synchronous events which could be involved in homeostatic regulation of neuronal assemblies, according to studies in rodent thalamic *ex vivo* slice preparations (Parri et al., 2001), rodent hippocampus slices (Bezzi et al., 2004; Fellin et al., 2004), and *in vivo* mouse neocortex (Poskanzer and Yuste, 2016). Furthermore, recent studies have suggested that conformational changes in the intracellular c-terminal domain of NMDAR may induce metabotropic-like signaling events in the absence of ionotropic functions, for example inducing long-term depression in rat hippocampal slices and cultures (Nabavi et al., 2013; Aow et al., 2015; Dore et al., 2015, 2016; Gray et al., 2016; Dore et al., 2017; Petit-Pedrol and Groc, 2021). Although the functions of extrasynaptic NMDAR-GluN2B in the primate dlPFC and SGC are completely unknown, our data suggest that the events such as those listed above could be more prevalent or frequent in the SGC than the dlPFC.

### Implications for vulnerability to depression

The SGC is overactive in patients with depression, and is a focus of deep brain stimulation for treating patients with intractable depressive symptoms (Mayberg et al., 2005). The dlPFC provides top-regulation of emotion through indirect projections to the SGC (Joyce et al., 2020; Arnsten et al., 2023), and symptoms of depression correlate with synapse loss from the dlPFC (Holmes et al., 2019). Given the extensive outputs of the SGC to brainstem and limbic areas, SGC overactivity may have outsized effects on brain states governing emotion and internal states (Hamilton et al., 2015; Arnsten et al., 2023). The dense expression of NMDARs in the SGC (Palomero-Gallagher et al., 2009) has led to the speculation that NMDAR antagonists such as ketamine and esketamine may have anti-depressant actions by quieting SGC output, similar to that produced by deep brain stimulation (Opler et al., 2016; Arnsten et al., 2023), especially as ketamine can normalize activity between dlPFC and SGC in marmosets (Alexander et al., 2021). One hypothesis is that NMDAR antagonism is enough to dampen SGC hyperactivity, and thus re-balance prefrontal networks and allow top-down regulation to resume (Joormann and Stanton, 2016; Opler et al., 2016; Yang et al., 2021; Arnsten et al., 2023).

It is possible that the extrasynaptic NMDAR-GluN2B in the primate SGC seen in the current study play a role in depression through their interactions with astrocytes, as schematized in <u>Figure 7</u>. The NMDAR has high affinity for glutamate, and it can be engaged at low glutamate concentrations, such as those observed in extracellular space (Sah et al., 1989; Paoletti et al., 2013). Astrocytes tightly control the concentration of extracellular glutamate via a number of mechanisms (Cuellar-Santoyo et al., 2022), such as glutamate uptake via the excitatory amino acid transporters (EAATs) (Todd and Hardingham, 2020), glutamate extrusion [e.g., via the cysteine-glutamate antiporter (De Bundel et al., 2011; Lewerenz et al., 2013; Soria et al., 2014)], or even vesicular glutamate release (Bezzi et al., 2004; de Ceglia et al., 2023). In mouse medial PFC (mPFC), extrasynaptic NMDAR-GluN2B are stimulated by a low tonic activation from ambient glutamate, and blocking that tonic NMDAR-GluN2B current abolishes depression-like behaviors, while preventing glial uptake of glutamate increases this tonic current (Miller et al., 2014). This suggests that extrasynaptic NMDAR are in equilibrium with glial glutamate transporters, and that this balance may govern aspects of mood-like behaviors in mice. Glial pathology has been reported in SGC and other PFC areas in postmortem brains of patients diagnosed with major depression [(Ongur et al., 1998); reviewed in (Cotter et al., 2001; Rajkowska and Stockmeier, 2013; Haroon et al., 2017; Banasr et al., 2021)], as well as in preclinical models [reviewed in (Miller et al., 2016)]. Some studies have found depression-related downregulation of the

**Figure 7.**
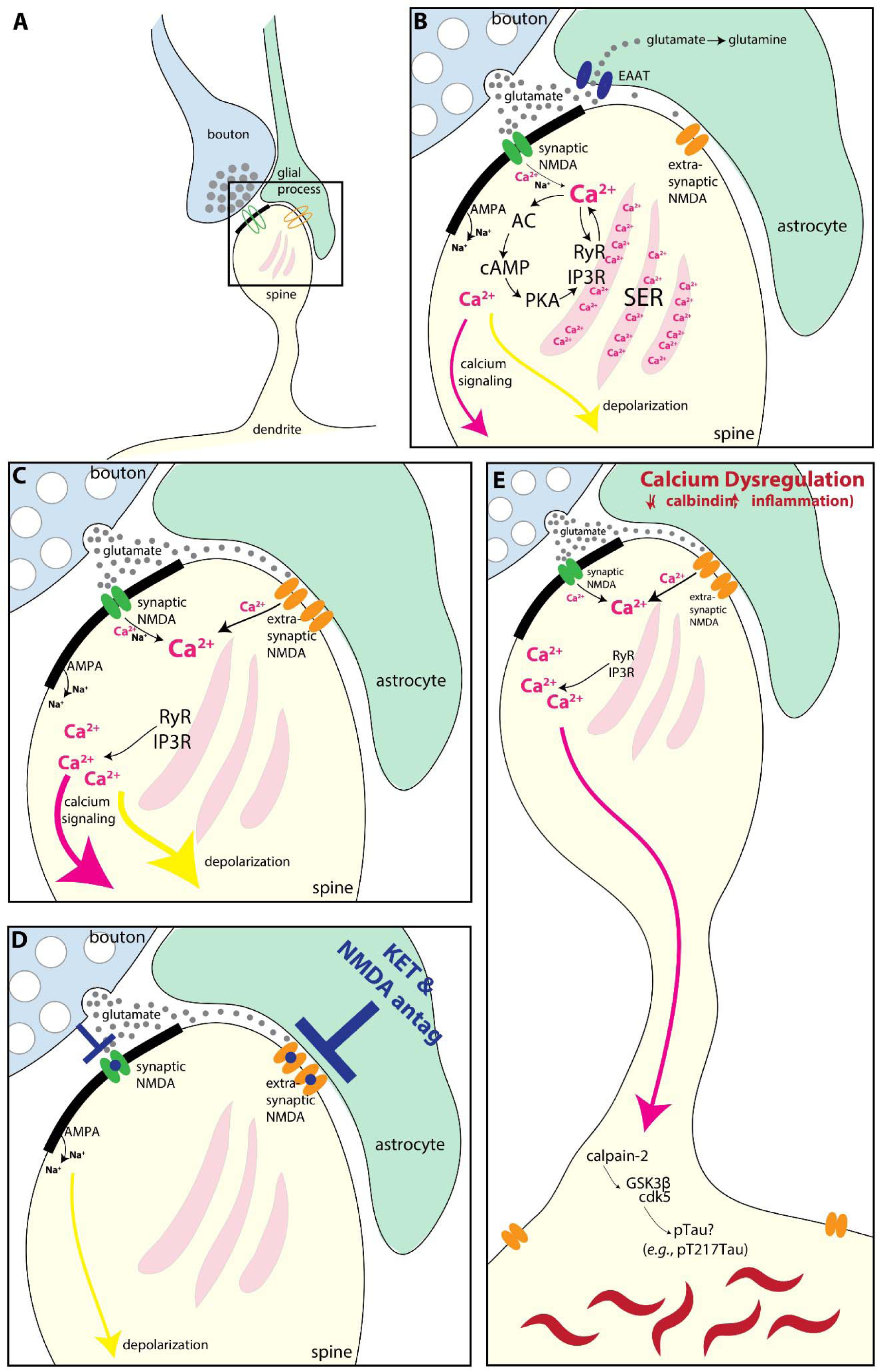
Schematic illustrating how extrasynaptic NMDAR-GluN2B may contribute to A25 hyperactivity, and/or calcium-mediated neurodegeneration-related events. ***A,*** Schematic depicting a spine (yellow), receiving a synapse from a glutamatergic bouton (blue), with an astrocytic leaflet (green) near the synapse. A synaptic NMDAR-GluN2B is present in the synapse (green), and an extrasynaptic NMDAR-GluN2B (orange) is present near the astrocytic process. ***B,*** A more magnified view showing that the bouton releases glutamate (grey circles) toward the postsynaptic density (thick black band), where the green synaptic NMDAR-GluN2B is engaged and calcium ions (pink), as well as sodium (Na+) ions, flow into the spine. AMPA receptors in the post-synaptic density also allow influx of sodium ions (Na+), though these are emphasized less as they are not the focus of the current work. Incoming calcium ions can trigger feedforward calcium release(Berridge, 1998; Arnsten et al., 2021b), via 1) direct calcium-mediated calcium release from the smooth endoplasmic reticulum (SER) via activation of primarily Ryanodine receptors (RyR), and 2) by cAMP magnification of calcium release, whereby calcium activates AC to produce cAMP which activates PKA signaling. PKA in turn phosphorylates the SER calcium channels RyR and IP3R to further increase calcium release. Glutamate escaping out of the synaptic cleft is sequestered into the astrocyte via the EAAT, where it can be converted to glutamine. In the dlPFC, calcium influx can also lead to a reduction in delay-related firing via SK3 channels (Datta et al., 2024) (not shown). ***C,*** If the EAATs are perturbed or downregulated, then glutamate can more readily engage extrasynaptic NMDA-GluN2B. Evidence suggests that there may be glial pathology in A25 (i.e. the SGC) during states of depression [(Ongur et al., 1998) reviewed in (Cotter et al., 2001; Rajkowska and Stockmeier, 2013; Haroon et al., 2017; Banasr et al., 2021)]. Given the prevalence of extrasynaptic NMDA-GluN2B we have found in the present study, we hypothesize that these may contribute to A25 hyperactivity observed in depression, perhaps by engaging feedforward calcium mechanisms and increasing depolarization (yellow arrow). ***D***, Rapid acting antidepressants that antagonize NMDA receptors may work in part by blocking extrasynaptic NMDAR-GluN2B in the SGC (Miller et al., 2016). ***E,*** Calcium is normally tightly regulated by cytosolic buffers (like calbindin and mitochondria), and by phosphodiesterases (which catabolize cAMP). Loss of this regulation with aging and/or inflammation(Arnsten et al., 2021a; Arnsten et al., 2021b; Joyce et al., 2024b) can dysregulate feedforward calcium signaling. Very high levels of cytosolic calcium can activate calpain-2, which cleaves and disinhibits GSK3β and cdk5, kinases that hyperphosphorylate tau, producing toxic species like pT217Tau (Arnsten and Baudry, 2023). Further post-translational modifications lead to tau fibrillation and the formation of neurofibrillary tangles. AC, adenylyl cyclase; cAMP, cyclic adenosine monophosphate; cdk5, cyclin-dependent kinase 5; EAAT, excitatory amino acid transporter; GSK3β, glycogen synthase kinase 3 beta; IP3R, inositol triphosphate receptor; PKA, protein kinase A; SER, smooth endoplasmic reticulum spine apparatus; RyR, ryanodine receptor

EAATs (Todd and Hardingham, 2020) in the PFC (Miguel-Hidalgo et al., 2006) and SGC (Scifo et al., 2018), suggesting that they may be less able to regulate extracellular glutamate concentration. Increased extracellular glutamate in depression may engage extrasynaptic NMDAR (Rajkowska and Miguel-Hidalgo, 2007; Miller et al., 2016) to produce despair-like behavior (Miller et al., 2014), and may be a mechanism for SGC hyperactivity (Alexander et al., 2019b; Alexander et al., 2020). Another postmortem study found that the SGC of patients with depression featured an upregulation of quinolinic acid (Steiner et al., 2011), a metabolite of tryptophan produced in glia that acts as an NMDAR agonist (Pedraz-Petrozzi et al., 2020), implying that glial-mediated dysfunction beyond the dysregulation of extracellular glutamate may also occur.

Extrasynaptic NMDAR may cause SGC hyperactivity through several mechanisms (summarized in <u>Fig. 7</u>), including the direct ion influx via tonic currents (Miller et al., 2014; Miller et al., 2016). In addition, depolarization via ion influx may also be augmented by calcium-mediated calcium release from the SER, which creates much larger amplitude changes in internal calcium concentration compared to extracellular calcium influx alone (Ross et al., 2005). Calcium can also drive cAMP signaling, which in turn can drive more internal calcium release (<u>Fig. 7</u>). These calcium mechanisms have been implicated in delay-related persistent firing observed in primate dlPFC (Arnsten et al., 2021b; Datta et al., 2024), and the induction of synaptic plasticity and tuning of memory fields in rodent hippocampus (O’Hare et al., 2022). In layer III of dlPFC, high levels of calcium reduce delay-related firing via the opening of SK3 channels (Datta et al., 2024), and it is not known if this also occurs in SGC. Thus, one possibility is that increased intracellular calcium decreases firing of layer III dlPFC neurons via SK channel opening, while increasing the firing of neurons in the SGC, maintaining the brain in a more emotion-dominated state.

Internal calcium release from the SER has been specifically implicated in the toxic effects of NMDAR-GluN2B signaling (Ruiz et al., 2009). Internal calcium release can also trigger protein kinase C, which mediates chronic stress-related spine and synapse loss in rat mPFC (Hains et al., 2009), and this may play a role in the etiology of depression-related neuronal atrophy. Postmortem studies have found that somatostatin in SGC is reduced in depression (Tripp et al., 2011; Seney et al., 2015), and somatostatin receptors inhibit adenylyl cyclase and calcium channels [reviewed in (Casello et al., 2022)], dampening signaling pathways necessary for feedforward calcium release. Downregulated somatostatin signaling may represent the loss of a key regulatory element for internal calcium concentration in depressive patients, which may also contribute to hyperactivity.

Here, using MLIF and fluorescence intensity analysis, we have detected higher expression of GluN2B in pyramidal neurons than inhibitory neurons. In both the SGC and dlPFC, we found prominent extrasynaptic expression in the dendrites of putative inhibitory neurons of layer III in both SGC and dlPFC, with far lower synaptic expression, especially compared to the nearby spines and dendrites of putative pyramidal dendrites. When present in the synapse, NMDAR-GluN2B were often found only at the very edge of the PSD, suggesting that they do not participate prominently in evoked synaptic NMDAR transmission, consistent with findings from other studies [(Nyiri et al., 2003; Rotaru et al., 2011) reviewed in (Miller et al., 2016)]. In these synapses they could fill an ancillary role in second messenger synaptic signaling rather than contributing prominently to synaptic conductance. Like in pyramidal neurons, in inhibitory neurons extrasynaptic NMDAR mediate tonic currents in the absence of synaptic stimulation (Povysheva and Johnson, 2012; Riebe et al., 2016). The disinhibition hypothesis of antidepressant action by NMDA antagonists postulates that antagonism of NMDAR on inhibitory neurons provides disinhibition of pyramidal neurons, allowing for renewed plasticity and spine growth in pyramidal neurons (reviewed in (Miller et al., 2016; Zanos and Gould, 2018; Brown and Gould, 2024)). Our data could suggest that any NMDAR-GluN2B mediated actions in dlPFC and SGC layer III inhibitory dendrites may have an outsized effect on extrasynaptic NMDAR than synaptic NMDAR. Thus the disinhibition hypothesis may rely on i) inhibition of extrasynaptic NMDAR-GluN2B, or ii) inhibition of synaptic NMDAR mediated by other subunits, such as the NMDAR-GluN2C and –GluN2D, which may be more prominent in inhibitory neurons (Paoletti et al., 2013), and ketamine notably has higher affinity for these subunits (Kotermanski and Johnson, 2009; Khlestova et al., 2016).

The dlPFC undergoes volume loss and neuronal atrophy in depression, e.g., synapse loss and soma size decreases (Rajkowska et al., 1999; Grieve et al., 2013; Holmes et al., 2019), much like chronic stress-related atrophy found in rodent prelimbic cortex, which has some properties in common with dlPFC (Hains et al., 2009). Although the SGC also undergoes general volume loss in depression, (Drevets et al., 1997; Botteron et al., 2002), it is unclear how much of that can be attributed to glial atrophy (Ongur et al., 1998). Neurons in rat mPFC that project to the entorhinal cortex exhibit chronic stress-related atrophy, while those that project to the amygdala do not (Shansky et al., 2009), so effects in SGC may be circuit-specific as well. The ultrarapid effects of rapid-acting NMDAR antidepressants may perform an acute role in reducing A25 hyperactivity and removing the dominance of A25 in prefrontal networks supporting euthymia and internal states (Hamilton et al., 2015; Opler et al., 2016; Yang et al., 2021; Arnsten et al., 2023). Then, slightly slower effects mediated by antagonism of NMDAR on inhibitory neurons, and subsequent glutamate surge (Abdallah et al., 2018), or by antagonism of synaptic NMDAR activated by spontaneous vesicle release (Zanos and Gould, 2018) may contribute to renewed plasticity in neurons atrophied by chronic stress. These actions are likely critical in re-establishing balance for top-down regions that project to SGC, like the frontal pole, dlPFC and pregenual anterior cingulate cortex, that then more slowly reinvigorate spine regrowth and metaplasticity mechanisms across the PFC for sustained remission (Brown and Gould, 2024).

### Implications for vulnerability to neurodegenerative forces

Dysregulated calcium signaling has been implicated in the early events leading to sAD (Khachaturian, 1989, 1994; Arnsten et al., 2021c; Datta et al., 2021). Consistent with this hypothesis, the degree of *GRIN2B* expression, and *CALB1* (calbindin) expression in pyramidal cells across the cortical hierarchy roughly aligns with the pattern and sequence of tau pathology in sAD (Braak and Braak, 1991; Garcia-Cabezas et al., 2017; Burt et al., 2018; Yang et al., 2018; Joyce et al., 2020; Arnsten et al., 2021c; Arnsten et al., 2021b; Arnsten and Baudry, 2023). High levels of cytosolic calcium can activate calpain-2 (Wang et al., 2013b), which in turn activates GSK-3β and cdk5, the major kinases that hyperphosphorylate tau and exacerbate Aβ42 cleavage from APP [reviewed in (Arnsten and Baudry, 2023)], and amyloid-β toxicity (Kessels et al., 2013; Talantova et al., 2013; Tamburri et al., 2013; Birnbaum et al., 2015). A recent analysis found a loss of synaptic NMDAR-GluN2B and an increase in extrasynaptic NMDAR-GluN2B in the dlPFC of patients with sAD, in line with previous hypotheses that extrasynaptic NMDAR-GluN2B may play a role in sAD etiology (Escamilla et al., 2024). Given that depression is a risk factor for later development of sAD (Ownby et al., 2006), extrasynaptic NMDAR-GluN2B in the SGC may represent a vulnerable nexus for pathology and mood disorder in both depression and sAD.

Memantine is FDA-approved to treat moderate-to-severe sAD, and is a non-competitive antagonist and open channel blocker of the NMDAR (Kim et al., 2024). Memantine has at times been thought to target extrasynaptic NMDAR (Xia et al., 2010), although this is likely an oversimplification (Glasgow et al., 2017). The lack of benefit from memantine in early stages of sAD may be due to its antagonism at synaptic NMDAR in dlPFC and related brain circuits that are critical for cognitive processes, thus creating a mixed response profile when combined with inhibition of possibly detrimental extrasynaptic NMDAR signaling. By moderate to severe stages of sAD, enough synapse and spine loss may have occurred to attenuate the structural organization of dlPFC microcircuits, thus occluding the detriment due to antagonism of synaptic NMDAR-GluN2B in dlPFC. Memantine has only modest anti-depressant effects (Hsu et al., 2022), which may have to do with differential interactions based on subunit composition and desensitization state from nearby high internal calcium concentrations (Glasgow et al., 2017). Understanding the details of NMDAR subtype locations and physiological functions in the primate PFC and cingulate cortices may help to refine treatment strategies for mood and cognitive disorders.

## Funding

This research was funded by R01 MH130538-01A1 and NSF 2015276 to AFTA.

## Conflicts of interest

The authors have no conflicts of interest with this study.

## Acknowledgements

We thank Lisa Ciavarella, Tracy Sadlon, Sam Johnson and Michelle Wilson for their invaluable contributions.

---

**Figure S1.**
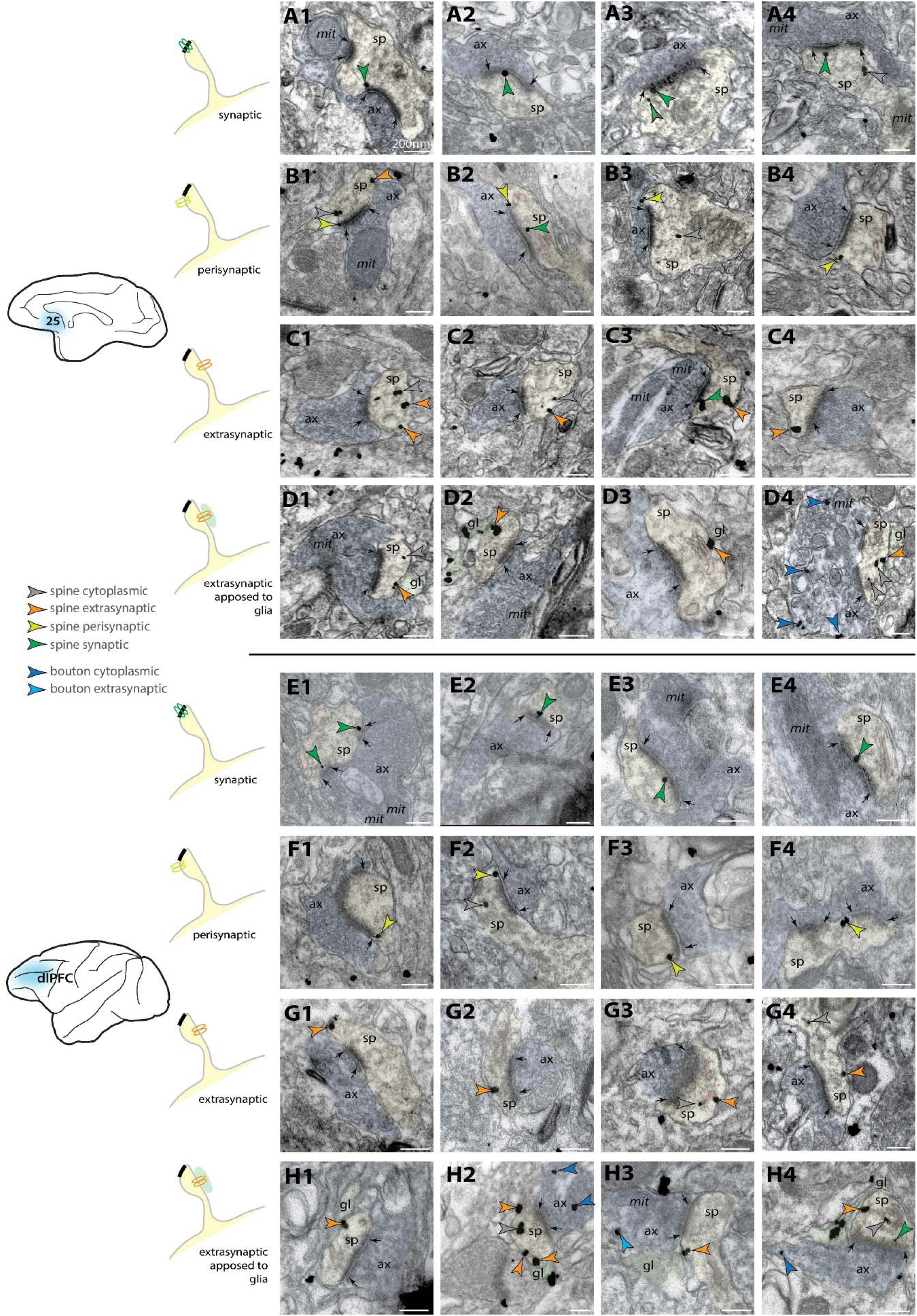
Additional examples of A25 and dlPFC NMDAR-GluN2B labeling in spines. ***A*,** Electron micrographs of A25 spines (pseudocolored yellow), receiving synapses (denoted by black arrows) formed by axon terminals (pseudocolored blue). NMDAR-GluN2B are prominent expressed in the synapse (green arrowheads), or at cytosolic locations (grey arrowheads). ***B***, A25 spines with prominent perisynaptic NMDAR-GluN2B (yellow-green arrowheads). NMDAR-GluN2B were classified as perisynaptic when within ∼100nm of membrane distance from the synapse. ***C,*** A25 spines with NMDAR-GluN2B expressed in the extrasynaptic membrane (orange arrowheads). ***D*,** A25 spines with extrasynaptic NMDAR-GluN2B apposed to glial-like processes (pseudocolored green), which sometimes also express NMDAR-GluN2B (*e.g.*, **D2**). Presynaptic cytosolic NMDAR-GluN2B (blue arrowheads) are occasionally evident (*e.g.*, **D4**). ***E-H,*** same as above but for dlPFC spines. Scale bars, 200nm; ax, axon; mit, mitochondria; sp, spine

**Figure S2.**
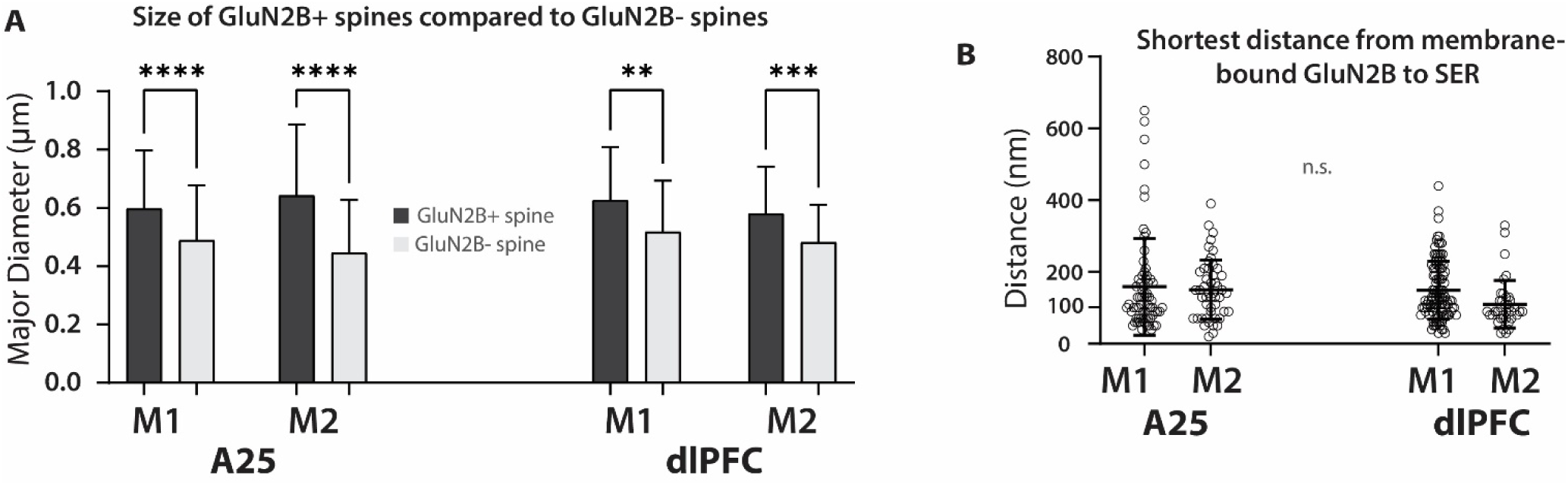
Quantitative characteristics of NMDAR-GluN2B+ spines and proximity of NMDAR-GluN2B to SER in spines. ***A,*** Plot of the major Feret’s diameter of spines in A25 and dlPFC that were GluN2B+ and GluN2B- in plane. Error bars depict standard deviation. One-way ANOVA with post-hoc Tukey test, F(7,1343)=26.69, p<0.001. ***B***, Swarm plot depicting the shortest distance from membrane bearing the NMDAR-GluN2B to the nearest spine apparatus SER membrane for all membrane-bound GluN2B immunogold particles found in spines. Swarm plot also depicts mean and standard deviation. *, p < 0.05; **, p< 0.01, *** p< 0.001; ****, p<0.0001;

**Figure S3.**
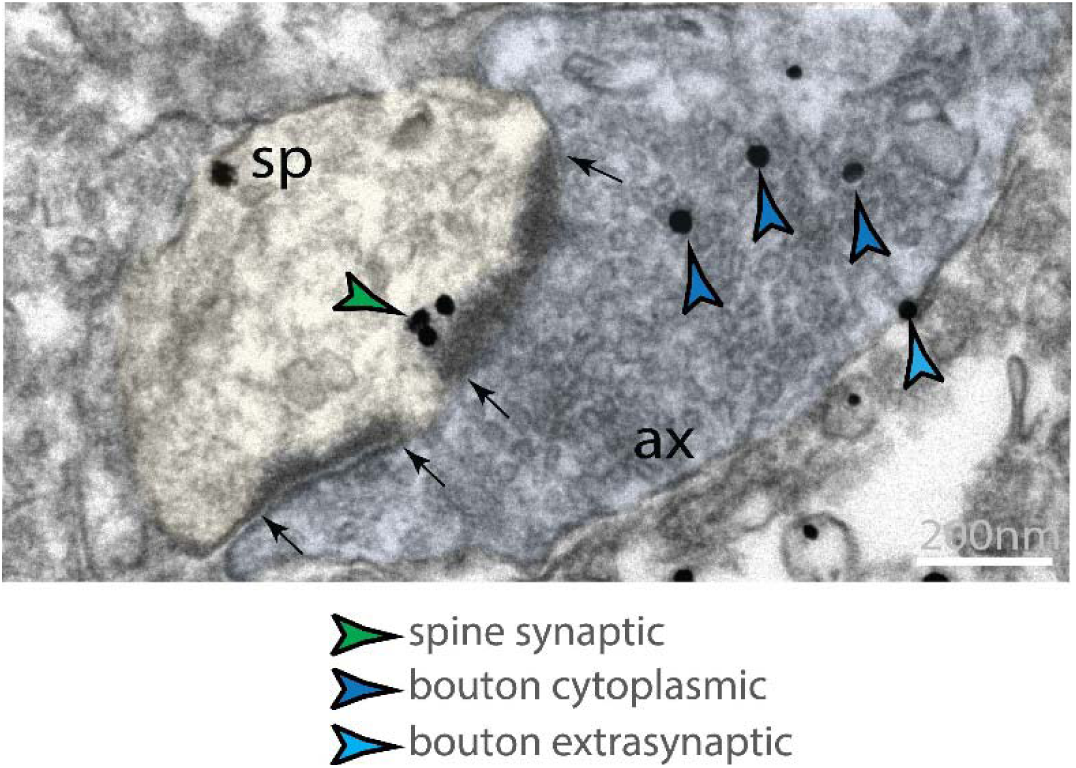
Presynaptic NMDAR-GluN2B. An A25 spine (sp, pseudocolored yellow) with prominent NMAR-GluN2B synaptic labeling (green arrowhead), and presynaptic NMDAR-GluN2B in the bouton (ax, pseudocolored blue) in the cytosol (darker blue arrowheads) amidst the vesicles, or on the extrasynaptic bouton membrane (lighter blue arrowhead). Black arrows denote the boundaries of the perforated synapse.

**Figure S4.**
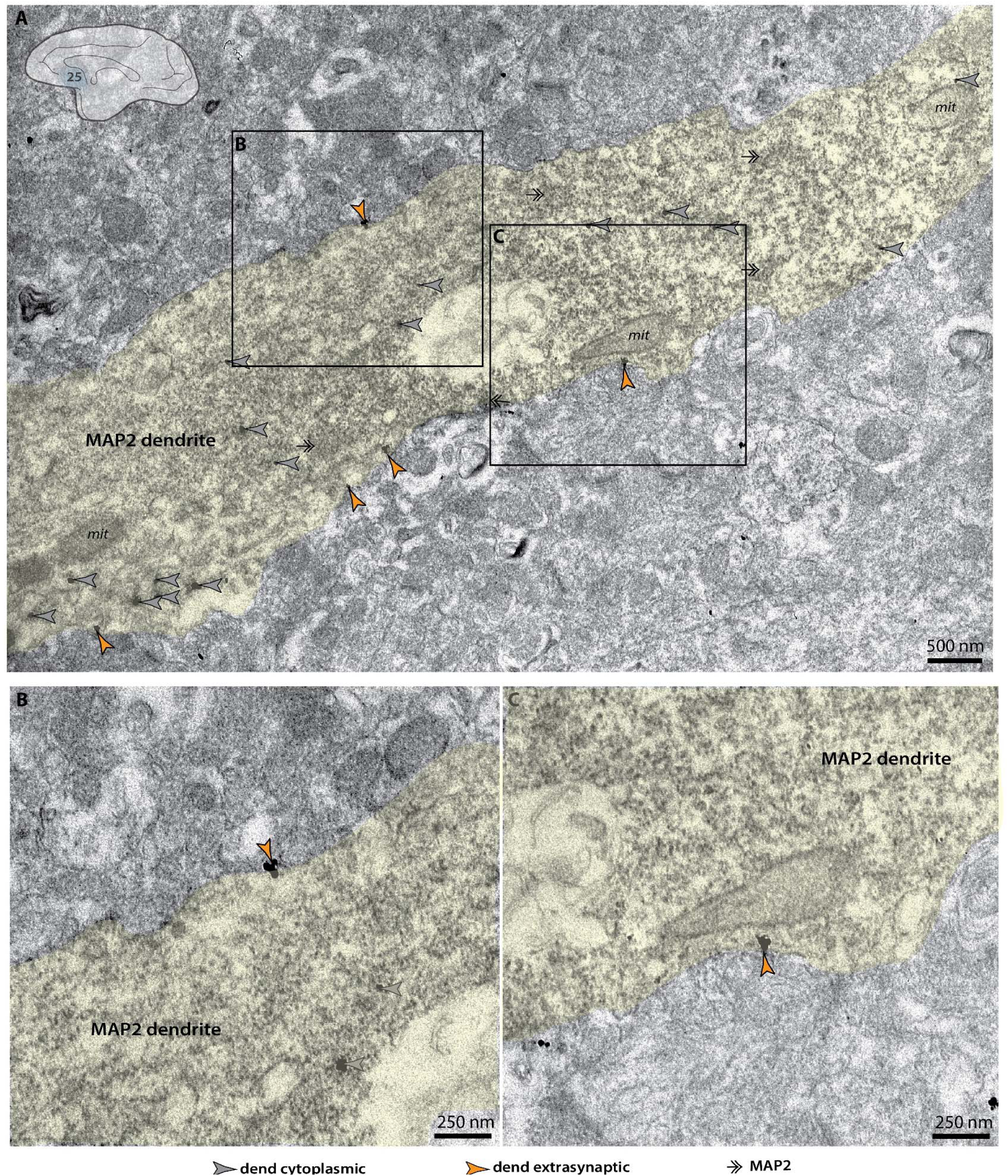
Extrasynaptic NMDAR-GluN2B in MAP2+ dendrite in A25. ***A***, A MAP2+ putative excitatory dendrite (pseudocolored yellow) with NMDAR-GluN2B labeling at intracellular (grey arrowheads) and extrasynaptic (orange arrowheads) locations. ***B,C***, Insets from **A**.

**Figure S5.**
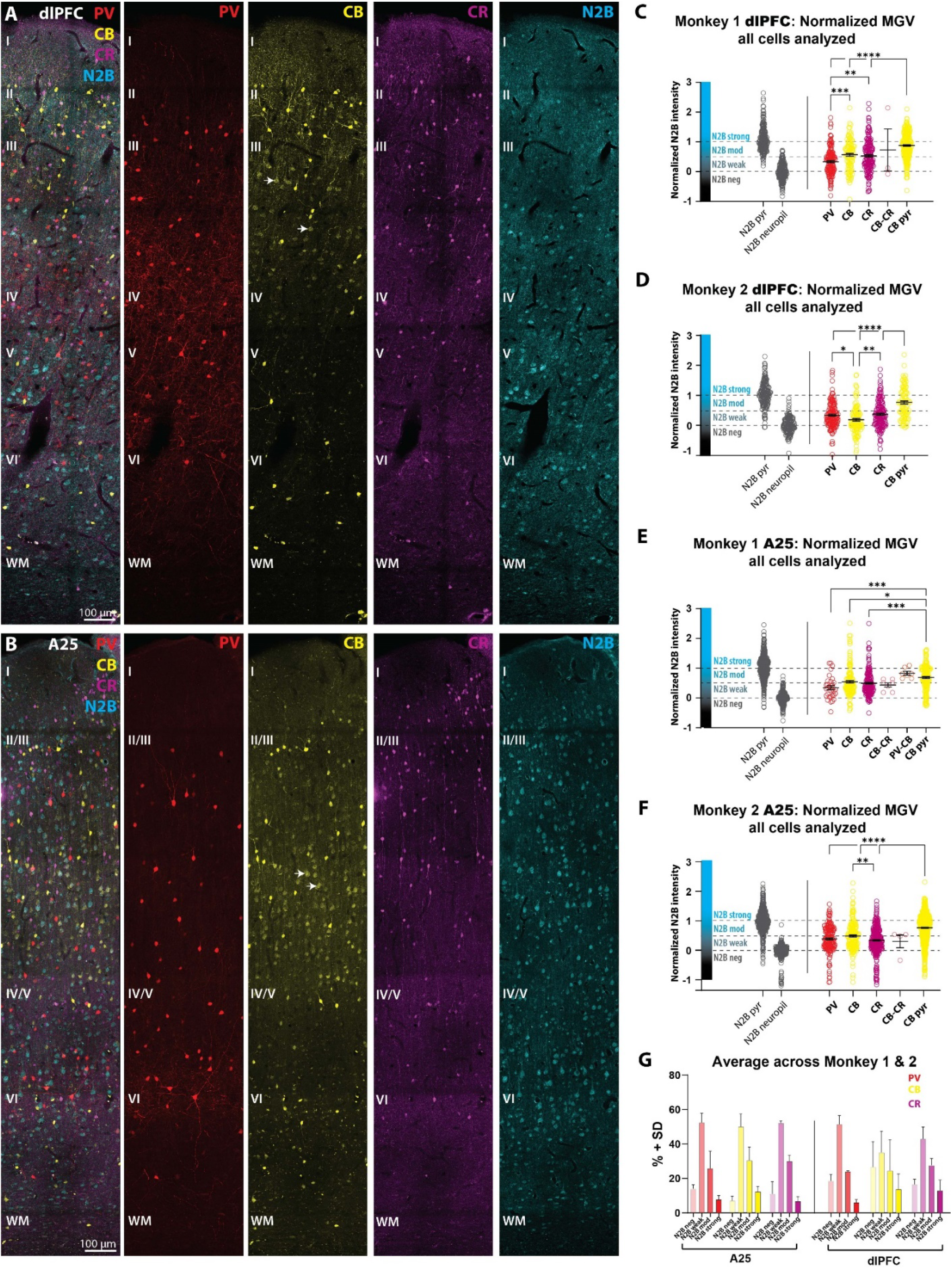
Supplemental MLIF and individual analyses for A25 and dlPFC NMDAR-GluN2B expression across CBP types in Monkey 1 and Monkey 2. Tiled images obtained via confocal microscopy of immunolabeled PV (red), CB (yellow), CR (magenta), and NMAR-GluN2B (cyan) in A25 (**A**) and dlPFC (**B**) across all laminar compartments. Images were systematically sampled from layer III. For each image, we isolated the NMDAR-GluN2B channel. Then we segmented i) NMDAR-GluN2B+ pyramidal-like neurons, as judged by morphology; and ii) manually selected immunonegative regions of tissue with no labeled NMDAR-GluN2B processes. We measured the mean intensity (MI) in all of these traces, and took the mean for each category (GluN2B+ pyramidal neurons or immunonegative background). We then used the other channels to segment the somata of PV, CB, and CR neurons. Then the MI was measured for each PV, CB, and CR neuron. We then used the mean NMDAR-GluN2B expression of the pyramidal neurons and of the immunonegative “neuropil” regions to create a normalized index of expression for each inhibitory neuron, where 0 was the average across sampled neuropil regions, and 1 was the average across sampled pyramidal-like neurons. This index forms the y-axis for **C-F**. Individual circles for each plot represent a cell, and these data were pooled across images after normalization. We divided the index into four equal bins from [0,1], delineated by dotted lines (Negative, at or below the average MI across sampled immunonegative regions; Weak; Moderate; or Strong, which was defined as at or above the average MI across sampled pyramidal neurons). Compiled data for Monkey 1 in A25 (**C**, One-way ANOVA, F(5,562)=6.314, p<0.0001, with post-hoc Tukey test) and dlPFC (**D** One-way ANOVA, F(4,701)=35.97, p<0.0001, with post-hoc Tukey test), and for Monkey 2 in A25 (**E,** One-way ANOVA, F(4,1693)=75.90, p<0.0001, with post-hoc Tukey test) and dlPFC (**F,** One-way ANOVA, F(3,542)=30.23, p<0.001, with post-hoc Tukey test)*. **G,*** Mean percent across cases of CBP+ inhibitory neurons by type that fell into Negative, Weak, Moderate, or Strong bins. CB, calbindin; CBP, calcium-binding protein; CR, calretinin; PV, parvalbumin; WM, white matter. *, p < 0.05; **, p< 0.01, *** p< 0.001; ****, p<0.0001

**Figure S6.**
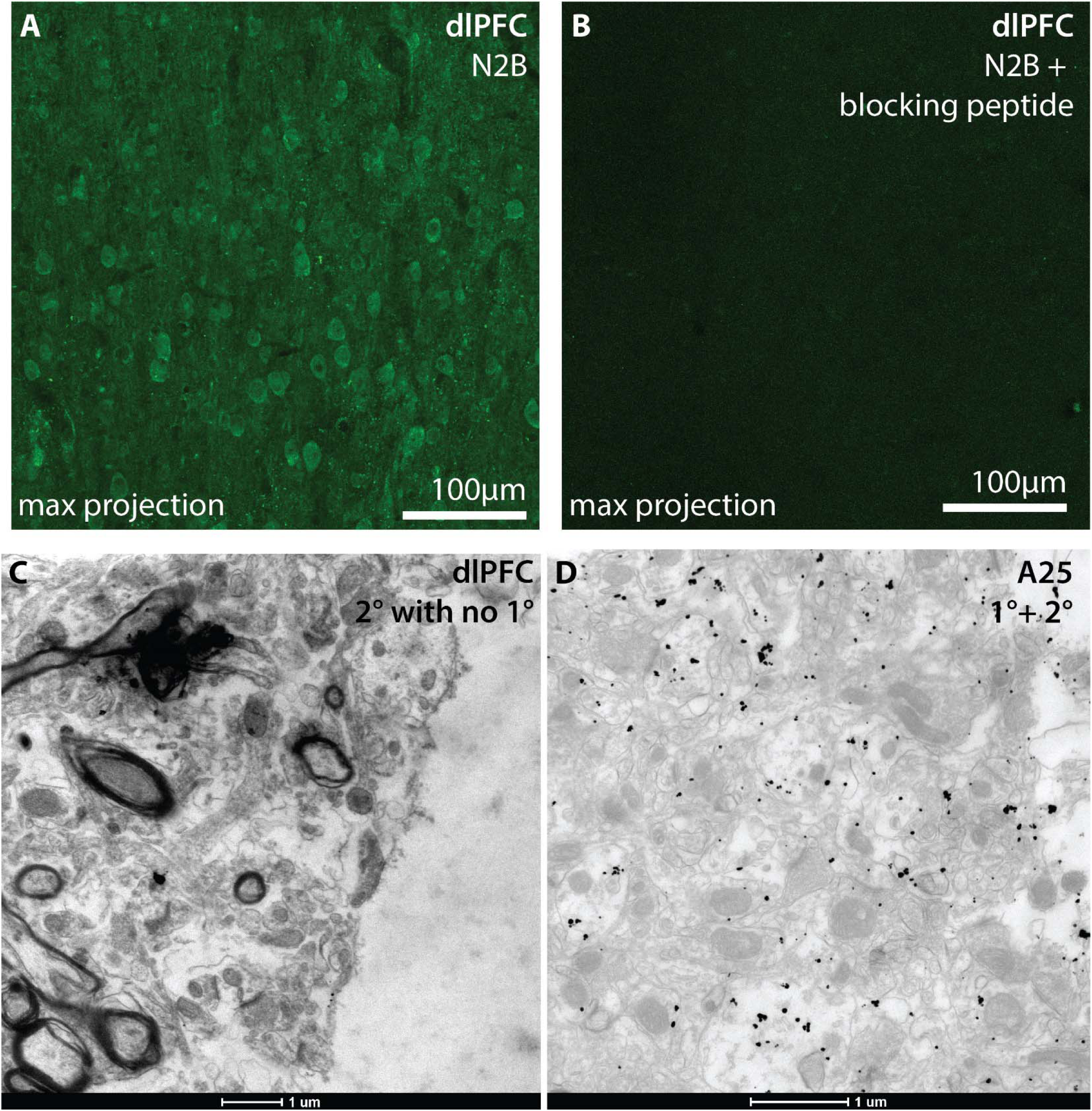
Immunohistochemical controls. ***A***, Confocal microscopy fluorescent images depicting layer III dlPFC single-labeled for NMDAR-GluN2B using the Alomone rabbit anti-GluN2B (cat #AGC-003) antibody. ***B***, A control section incubated in a separate aliquot of primary antibody solution used in **A**, but with the addition of the Alomone blocking peptide (cat #BLP-GC003). The negligible labeling observed after imaging with the same parameters as **A** suggests that the paratope region of the antibody has very low non-specific interactions in our tissue. ***C***, Electron micrograph from a control section of tissue with the omission of the primary antibody (1°) but all other procedures intact. The image was captured at the edge of the tissue “coming into plane”, where antibody penetration is at its greatest, often producing some noise or background level labeling. ***D***, Electron micrograph from the same batch of tissue, from a section treated with both the primary antibody and secondary antibody (1°), and all other procedures held constant. The image is taken near the edge of the tissue “coming into plane” (right), where antibody penetration is typically greatest, often producing some noise or background level labeling.

## Notes

### Competing Interest Statement

The authors have declared no competing interest.

